# Trace impurities in test stimuli can seriously compromise chemosensory studies

**DOI:** 10.1101/503169

**Authors:** Dirk Louis P. Schorkopf, Béla Péter Molnár, Marit Solum, Mattias C. Larsson, Jocelyn G. Millar, Zsolt Kárpáti, Teun Dekker

## Abstract

The discovery of olfactory receptors and major technological advances have greatly accelerated our understanding of chemosensory mechanisms. However, some of this rapid progress may be compromised by inadequate knowledge or characterization of the purity of chemical stimuli used to challenge olfactory or other chemoreceptors when mapping their response profiles. Here, we provide strong evidence that the presence of trace impurities in test stimuli can completely obscure true ligand-receptor relationships. DmOR7a, an olfactory receptor of the vinegar fly (*Drosophila melanogaster*) has been reported to respond to several long-chain aliphatic ligands such as a putative *Drosophila* pheromone^1^, the pheromone of the silkworm moth *Bombyx mori^2^*, and a common fatty acid, linoleic acid^3^. By contrast, we show that DmOR7a responds with high sensitivity to volatile impurities and degradation products present in minute quantities in authentic standards of those compounds, but not to the standards themselves. Responses to impurities can easily go unnoticed due to two main factors. First, the sensitivity of receptors to key ligands may be greater than that of analytical chemistry instruments used to check sample purity. Second, the concentration of highly volatile impurities in an odour puff may be orders of magnitude higher than the main component of a sample, due to the large differences in vapour pressures between the impurities and the main component. Issues concerning impurities are not limited to studies on olfaction that use odour puffs to characterize receptor-ligand interactions, but may affect all studies on chemosensation, from molecular biology and in-silico predictions to behaviour. Purity, which is crucial in receptor-ligand studies, is always implied, but rarely checked rigorously. To avoid misinterpretations, a proper account of all compounds present in test stimuli and an unequivocal confirmation of ligand affinity should accompany chemosensory studies.

---

The field of chemosensory sciences has progressed rapidly through molecular, genetic, and neurophysiological advances that permit the unravelling of the full sequence from perireceptor events to receptor-induced intracellular responses, downstream neuronal signalling and processing in the brain, and ultimately behavioural output^4–8^. A commonplace assumption in these studies is that standards used as chemosensory stimuli are pure, or alternatively, that observed responses are the result of interactions between the receptor and the nominal authentic standard.

In routine evaluations of *D. melanogaster* olfactory receptor (OR) affinities, we observed that AB4a neurons housed in antennal basiconic sensilla responded differently than anticipated. This neuron and its endogenous receptor DmOR7a are reported to be broadly sensitive to short-chain six-carbon aldehydes, alcohols, and esters^9^, but also to longer chain compounds, such as the silk moth pheromone, bombykol ((10*E*,12*Z*)-10,12-hexadecadien-1-ol)^2,10^, the *Drosophila* cuticular hydrocarbon, (*Z*)-9-tricosene (Z9T)^1^, and linoleic acid (LLA)^3^. We found that cartridges loaded with synthetic bombykol, Z9T, or LLA quickly lost activity with repeated puffs (Extended Data Fig. 1c, and Extended Data Fig. 2c). This was unexpected, because these long-chain compounds have low vapour pressures and would be expected to deliver a relatively constant stimulus dose over numerous puff cycles^11–13^. Indeed, such declines were not observed (Extended Data Fig. 1a,b) when using the same protocol to stimulate the pheromone receptor of *B. mori*, BmOR1^14^ (exogenously expressed in *D. melanogaster* T1 neurons, T1_BmOR1_) with bombykol, or when stimulating wildtype T1 neurons (expressing its cognate receptor DmOR67d^15^) with its ligand, the long-chain *Drosophila* pheromone *cis-*vaccenyl acetate ((*Z*)-11-octadecenyl acetate; *c*VA).

Furthermore, AB4a neurons responded equally well to bombykol on a filter paper or in paraffin oil^11^ (Extended Data Fig. 3a). This was counterintuitive, because non-volatile paraffin oil should retain bombykol, a long-chain aliphatic compound, and significantly reduce volatilization and hence stimulus intensity compared to bombykol applied to filter paper^11,13^. Indeed, responses of antennal trichoid T1 (sensitive to *c*VA) and T1_BmOR1_ neurons (sensitive to bombykol) were significantly attenuated when stimulated with air puffed over dilutions of *c*VA or bombykol dissolved in paraffin oil versus on filter paper (Extended Data Fig. 3b, c).

This cast doubt on whether the above-mentioned compounds were indeed ligands for AB4a neurons. To more rigorously test this, we used coupled gas chromatography-electroantennographic detection (GC-EAD), which separates the injected sample into its individual components and sequentially passes these over the antennal preparation. Thus, each antennal response can be unequivocally attributed to a defined peak, which generally represents a single pure compound. We found that the cleanly separated bombykol peak did not induce antennal depolarization in wildtype fly antennae (Fig. 1a; Extended Data Fig. 4 and 5), nor did bombykal (another reported ligand for AB4a neurons)^2,10^, Z9T, or LLA (Fig. 1b, Extended Data Fig. 5). The GC-EAD setup was clearly functioning properly because antennae of male *B. mori* responded strongly to bombykol (Fig. 1c), as did antennae of *D. melanogaster* expressing the bombykol receptor BmOR1 in T1 neurons (Fig. 1a). Wildtype fly antennae also responded as expected to *c*VA (Fig. 1, Extended Data Fig. 4).

**Figure 1.**
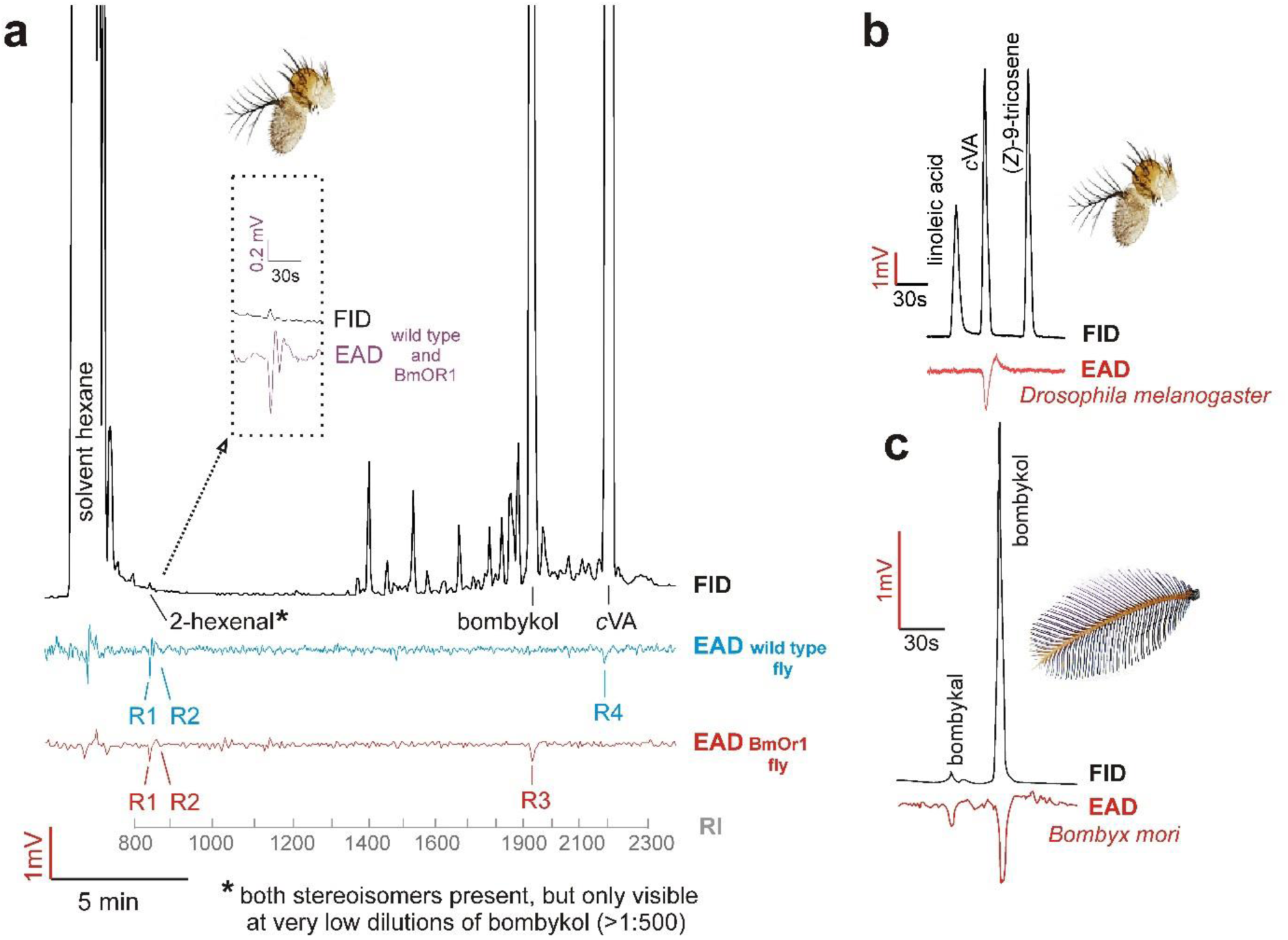
Representative responses of whole mount fly and moth antennal preparations to compounds eluting from a GC (GC-EAD) following injection of hexane solutions of synthetic standards. **A**, Wildtype fly antennae did not respond to the main component bombykol (n=30; shown is the average of n = 8 flies), but responded (R1, R2; see magnification of the trace) to volatile impurities eluting earlier (later identified as (*Z*)- and (*E*)-2-hexenal, see Table 1) as well as to the co-injected aggregation pheromone *c*VA (response R4). Replacing the *c*VA receptor (DmOr67d) with the receptor (BmOR1) substituted the response (average of n=8 flies) to *c*VA with a response to bombykol (R3). **B**, Wildtype fly antennae responded (n=5 flies) to *c*VA but not to LLA or Z9T in a mixed standard of LLA, *c*VA, and Z9T. **C**, Antennae of male *B. mori* responded to bombykol and bombykal (n=5).

Whereas bombykol did not elicit responses from wildtype antennae, responses were elicited by several impurities that eluted much earlier than bombykol (Fig. 1, R1 and R2). Using coupled GC-single-sensillum recordings (GC-SSR), we challenged AB4 sensillum preparations sequentially with these impurities. The above two low-molecular-weight impurities (R1 and R2) in the bombykol and bombykal samples induced strong responses in AB4a neurons (Fig. 2a). Coupled GC-mass spectrometry subsequently identified these as (*E*)-2-hexenal (E2H) and (*Z*)-2-hexenal (Z2H). E2H is a known ligand for AB4a neurons and their cognate receptor DmOR7a^9^. Puffs from cartridges loaded with 2-hexenal in amounts corresponding to those in our bombykol sample induced antennal responses comparable to those seen with our bombykol sample (Fig. 2b). We further tested the role of 2-hexenal by removing E2H and Z2H from the sample, predicting that this would significantly reduce the responses of AB4a neurons. Thus, reduction of the sample with sodium borohydride, which reduces E2H and Z2H to the far less stimulatory^9^ alcohols (*E*)- and (*Z*)-2-hexenol, dramatically attenuated the responses of AB4a neurons, as did reduction of synthetic E2H itself (Fig. 3a, b). E2H may arise from oxidative degradation of bombykol and bombykal at carbon 10, similar to oxidative degradation of unsaturated fatty acids^16^. Interestingly, a freshly synthesized batch of bombykol obtained from the same company contained significantly less E2H than the original sample, and accordingly induced lower responses from AB4a neurons (Extended Data Fig. 2c, 6).

**Figure 2.**
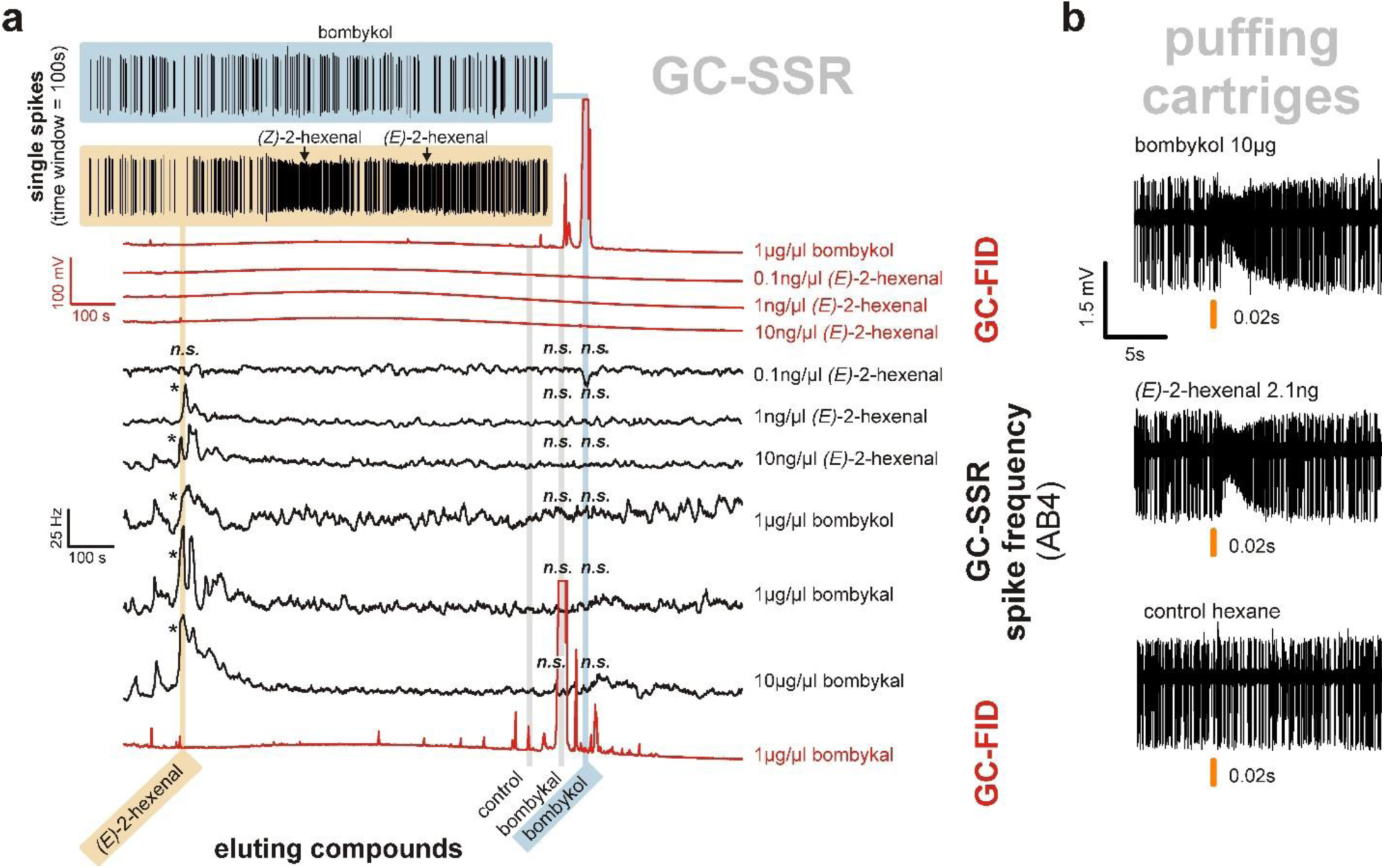
Responses of *Drosophila* antennal basiconic 4a neurons to bombykol, bombykal, and E2H. **A**, Responses of AB4a neurons to chromatographically separated dilutions of synthetic standards of bombykol, E2H, and bombykal. Whereas the peaks from pure bombykol or bombykal never induced a response (top blue trace), E2H did (yellow example trace), at all concentrations above 0.1 ng/µl. In red: GC-FID traces, in black: responses of AB4A neuron converted to Hz. **B**, Single sensilla were treated with puffs from cartridges loaded with either bombykol or E2H (loaded with amounts equivalent to the amount of E2H impurity in the bombykol standard), eliciting comparable neuronal responses. Asterisks (*) indicate significant differences versus the control (n=7, p<0.05; Holm-Sidak multiple comparisons versus control) following One Way Repeated Measurements ANOVA. n.s. = not significant.

**Figure 3.**
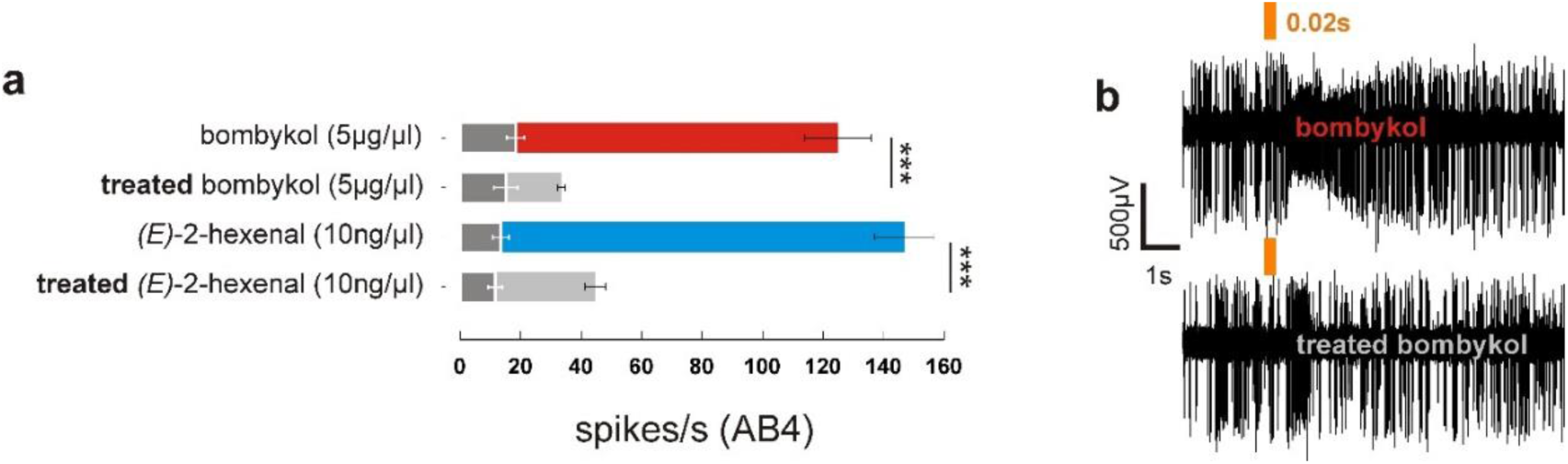
Chemically reducing C6 aldehydes to alcohols eliminated responses (means ±95%CI) of antennal basiconic 4a neurons that were typically elicted by the untreated synthetic bombykol and E2H (see also fig. 2B). **A**, Puffs with the synthetic standard after reduction with sodium borohydride, which converted E2H and Z2H to the corresponding alcohols, elicited dramatically attenuated responses to the reduced products. Reducing synthetic E2H under the same conditions gave analogous results. Overlaid dark grey bars with white whiskers show the results obtained for the pre-puffing control periods. Asterisks (***) indicate highly significant differences (paired t-test, n=8 different flies, p<0.001). **B**, Example traces of stimulation of the AB4a neuron with synthetic bombykol before (top) and after (bottom) sodium borohydride treatment.

We subsequently assessed whether samples of LLA and Z9T also contained E2H or other AB4a-stimulating impurities. Indeed, GC-SSR analyses of LLA and Z9T samples showed responses from AB4a neurons at the retention time of E2H (Extended Data Fig. 7), although weaker than bombykol, likely due to the substantially lower amounts of E2H in the samples (Extended Data Figs. 8). Similar to bombykol, Z9T samples from two different suppliers elicited markedly different response amplitudes from AB4a neurons (Extended Data Fig. 2a,b), suggesting that impurities, rather than Z9T itself, induced the responses. This is underscored by the fact that AB4a neurons responded more strongly to puffs of synthetic (*Z*)-7-tricosene (Z7T) than Z9T (Extended Data Fig. 2a), likely because this sample contained ~10-fold more E2H than either of the Z9T samples. Z7T is a male cuticular pheromone of *D. melanogaster*^17,18^ and present in much higher amounts than Z9T^18^, but was excluded in the electrophysiological evaluations of the above-mentioned study^1^. Finally, we assessed whether AB4a neurons responded to biological samples containing Z9T: odour puffs from a cartridge loaded with a cuticular extract from 350 (mixed sex) or 70 *Drosophila* (separated sexes) containing up to ~15 µg of Z9T, did not elicit significant responses (Extended Data Fig. 9). None of the above observations fit with a Z9T-mediated role for AB4a neurons in aggregation and oviposition^1^, but instead show that AB4a neurons respond to impurities in synthetic Z9T samples, rather than to Z9T itself.

In the above analyses, each of the synthetic samples contained approximately 5% impurities constituting numerous tiny peaks (e.g. Fig. 1, Fig. 4), and sensory neurons appeared extraordinarily sensitive to some of these trace impurities, even when below the GC detection threshold (~ 1 picogram^11,13^). Thus, standard GC analysis may not suffice for detection of confounding impurities in samples.

**Figure 4.**
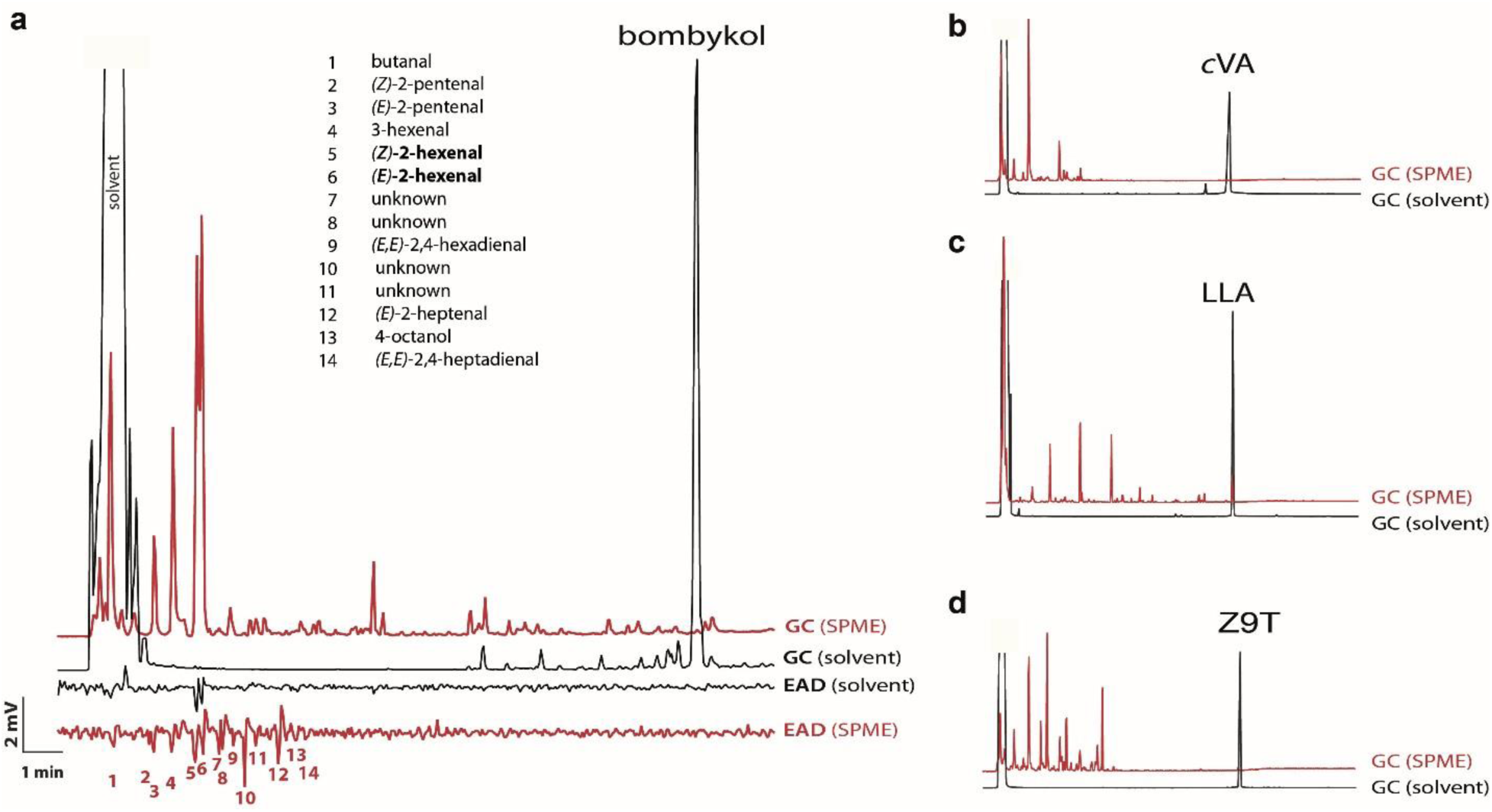
Concentrations of compounds in the vapour phase were orders of magnitude different than in the bulk liquid sample. **A**, Injections of solutions of standards (black) show very low abundance of impurities, but the concentrations of volatile impurities can be orders of magnitude greater in the headspace above the liquid samples (red, sampled using SPME), as illustrated with the synthetic standard of bombykol. Accordingly, antennae may respond to confounding impurities in headspace samples (red) much more intensely than when samples are injected as solutions (black), as exemplified with bombykol. **B**, **C** and **D** illustrate the same phenomenon with standards of *c*VA, LLA, and Z9T, in which the concentrations of volatile impurities in the headspace are magnified enormously relative to the main component, in comparison to the concentrations in the bulk samples.

The problem may be compounded when samples are puffed over antennal preparations. During puffing, the transition of compounds from the liquid to the vapour phase is largely dependent on their vapour pressures^11,13^. Accordingly, the relative proportions of compounds in the vapour may be massively different than their proportions in the liquid phase. For example, because the calculated vapour pressure of E2H at 25ºC is 629 Pa, versus 7.59 × 10^−4^ Pa for bombykol (Supplementary Table 5, Extended Data Fig. 10^19^), the headspace above a sample of bombykol containing 0.1% E2H would contain far more E2H than bombykol. Consequently, the composition of the bulk sample may be entirely unrepresentative of the composition of the headspace used for stimulation. Indeed, the headspace of bombykol, Z9T, LLA and cVA, sampled with Solid Phase Microextraction (SPME)^20^, was dominated by numerous volatile impurities (Fig. 4, Extended Data and Supplementary Tables 1–4), whereas the main compound was barely detectable. Many impurities in the headspace of samples elicited consistent EAD responses (Fig. 4, Extended Data Tables 1–4), among which were several ligands for receptors other than DmOR7a^9^.

It is common practice in chemoreceptor studies to prepare panels of chemical species at fixed amounts or concentrations so to assign ligands to receptors, sensory neurons, processing networks and behaviour^9,11^. In addition to the potentially confounding impurities present in synthetic standards, we emphasize that the precise amount and ratio of molecules reaching the target might vastly differ from the prepared/intended amount^11^ if factors such as different vapour pressures or solubility are neglected.

Importantly, impurities can affect chemosensory studies even when a compound has been unequivocally linked to a target neuron. For instance, in our study, all the synthetic samples contained impurities that induced responses in sensory neuron types other than AB4a neurons (see Fig. 4a and Extended Tables 1- 4^9^). The above samples thus stimulated non-target sensory neurons with unknown effects on downstream neural integration and behavioural output^21^.

In the present study, a single neuron-receptor combination, AB4a-DmOR7a, served to illustrate how the extraordinary sensitivity of receptors to their key ligands, vast differences in vapour pressures between a putative test chemical and its impurities, or a combination thereof^11,22–24^ can skew or even completely confound the results of otherwise elegantly crafted studies. This issue of impurities is ubiquitous and pernicious, potentially affecting any study involving chemoreceptors and sensory neurons, and the correct interpretation of downstream neuronal outputs, signal integration in the brain, and finally behavioural responses. To minimize errors due to impurities and more reliably correlate ligands with their receptors, neural circuits and behaviour, we advocate using methods such as GC-EAD or GC-SSR which separate out impurities and deliver known amounts of pure ligands to their targets.

## Acknowledgements

We thank Prof. W. Francke for comments during data collection and writing, Prof. Walter Leal for BmOR1 *Drosophila* lines, and Prof. Barry Dickson for the DmOR67D-Gal4 *Drosophila* line. The research was supported by the following grants: ICE^3^ (a Linnaeus grant to the unit of Chemical Ecology, DLPS, MS, MCL, TD), Vetenskapsrådet (621-2014-4816, TD), TASENE (a grant for research collaboration with Tanzania, Tasene/Sida/2/2012, DLPS), Carl-Fredrik von Horns fund of the Royal Swedish Academy of Agriculture and Forestry (H11-0195-CFH-01, DLPS), Guest Scientists Fellowship Programme of the Hungarian Academy of Sciences (VK-003/2016, TD, ZK), Hungarian Scientific Research Fund - NKFIH (K 119844, ZK, BPM), National Research, Development, and Innovation Office (GINOP-2.3.2-15-2016- 00051, ZK, BPM).

## Author Contributions

Conceived the original idea of the significance of impurities in chemosensory studies (TD), initiated, conceptualized and coordinated the research (DLPS, TD), designed the experiments (DLPS, MS, MCL, BPM, ZK, TD), executed the experiments (DLPS, MS, MCL, BPM, JM, ZK, TD), analyzed chemical data and identified impurities (DLPS, BPM, JM), analyzed physiological data (DLPS, BPM, ZK, TD), performed statistics (DLPS), wrote initial versions of the manuscript (DLPS, TD), commented and improved the manuscript (DLPS, MS, MCL, BM, JM, ZK, TD).

**Extended data** is available. [to be filled in after acceptance for publication]

**Supplementary information** is available. [to be filled in after acceptance for publication]

**Reprints and permissions information** [to be filled in after acceptance for publication]

---

**Extended Data Figure 1.**
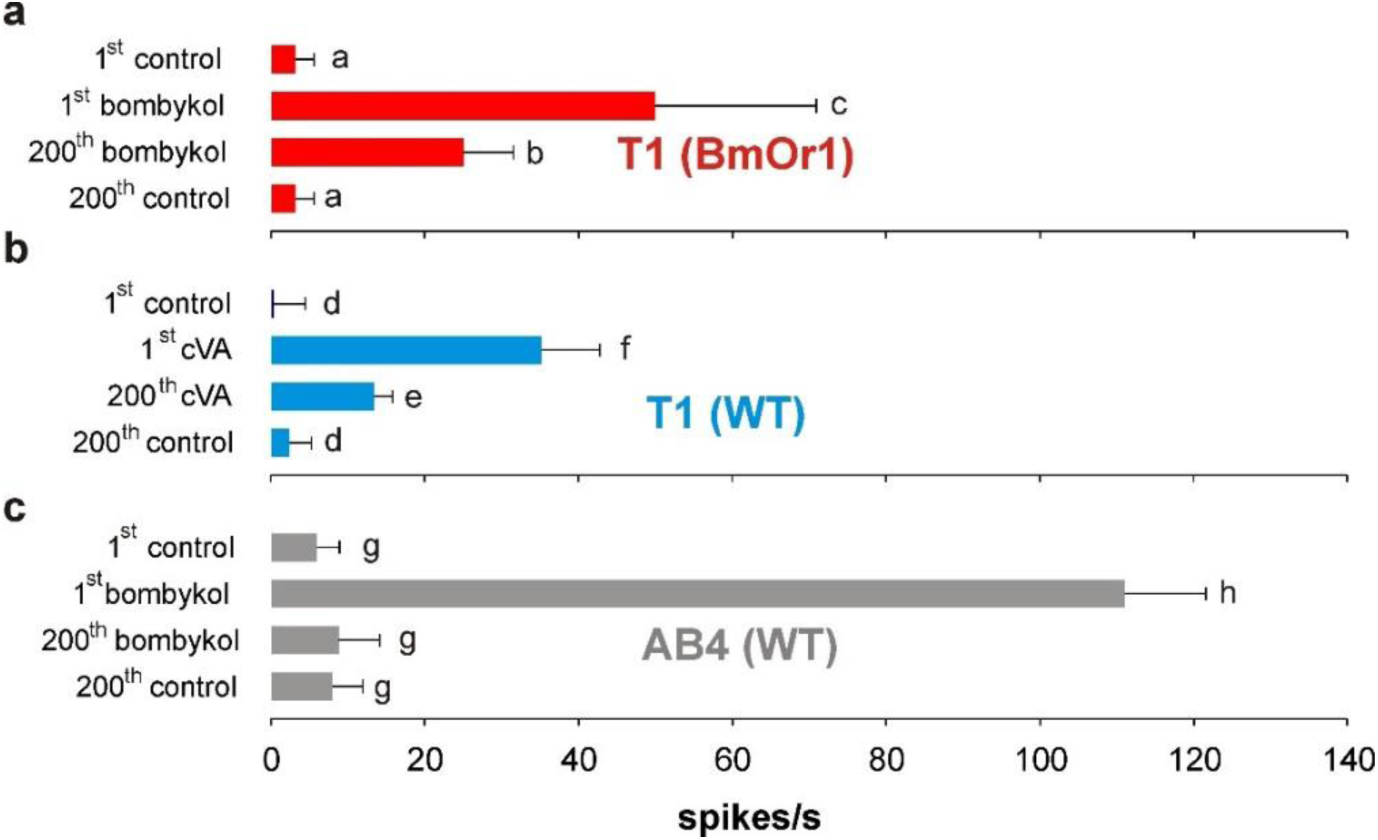
Responses (means +95%CI) of sensory neurons to stimulation from fresh stimulus cartridges versus cartridges after 200 puffs. While T1 neurons expressing BmOR1 (**A**) and wild type T1 neurons (B) responded to both fresh *c*VA-loaded cartridges or cartridges puffed 200 times in succession, AB4a neurons only consistently responded to fresh cartridges (**C**). Values with the same letters are not significantly different from each other (One Way Repeated Measurements ANOVA, followed by Holm-Sidak multiple comparisons; p>0.05, n=5 different flies).

**Extended Data Figure 2.**
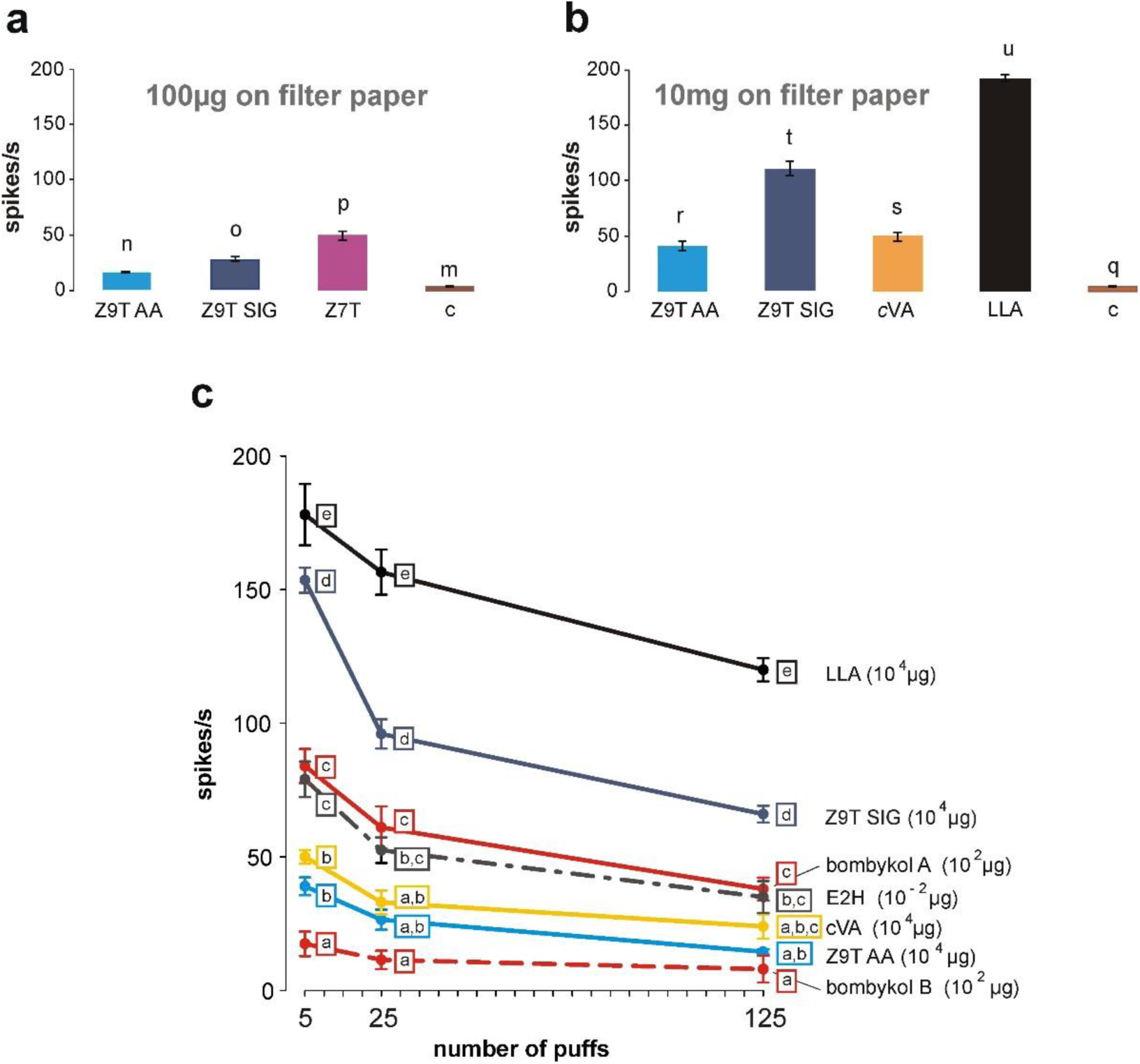
Single sensillum recordings (means ± SE) from AB4a neurons via puffing using different compounds, batches and depletion series. **A**, Z9T SIG (100 µg on filter paper) from Sigma-Aldrich induced stronger responses by puffing than Z9T AA from Alfa-Aesar. However, a synthetic sample of Z7T, which is more abundant than Z9T on *D. melanogaster* cuticle, induced stronger AB4a responses (n=7 flies) than either of the Z9T synthetic samples. **B**, This also held true at higher doses (10 mg on filter paper; n=7 flies). At these amounts, even a sample of cVA induced a response (n=4 flies). LLA was included as a reference and gave stronger responses (n=4 flies) than Z9T and cVA. **C**, responses of AB4a neurons (n=4 flies) to odour cartridges after 5, 25, or 125 repeated stimulations with E2H (10 ng on filter paper) or synthetic bombykol (100 µg on filter paper). Depletion of responses to Z9T (SIG, Sigma-Aldrich; AA, Alfa Aesar), LLA and cVA standards (10 mg on filter paper) with increasing numbers of puffs are also shown. A new batch of bombykol (dotted red line) induced significantly lower responses from AB4a neurons compared to a batch (solid red line) received 2 years earlier (see also Extended Data Figure 6). Values with the same letters are not significantly different from each other (One Way Repeated Measurements ANOVA, followed by Holm-Sidak multiple comparisons; p>0.05).

**Extended Data Figure 3.**
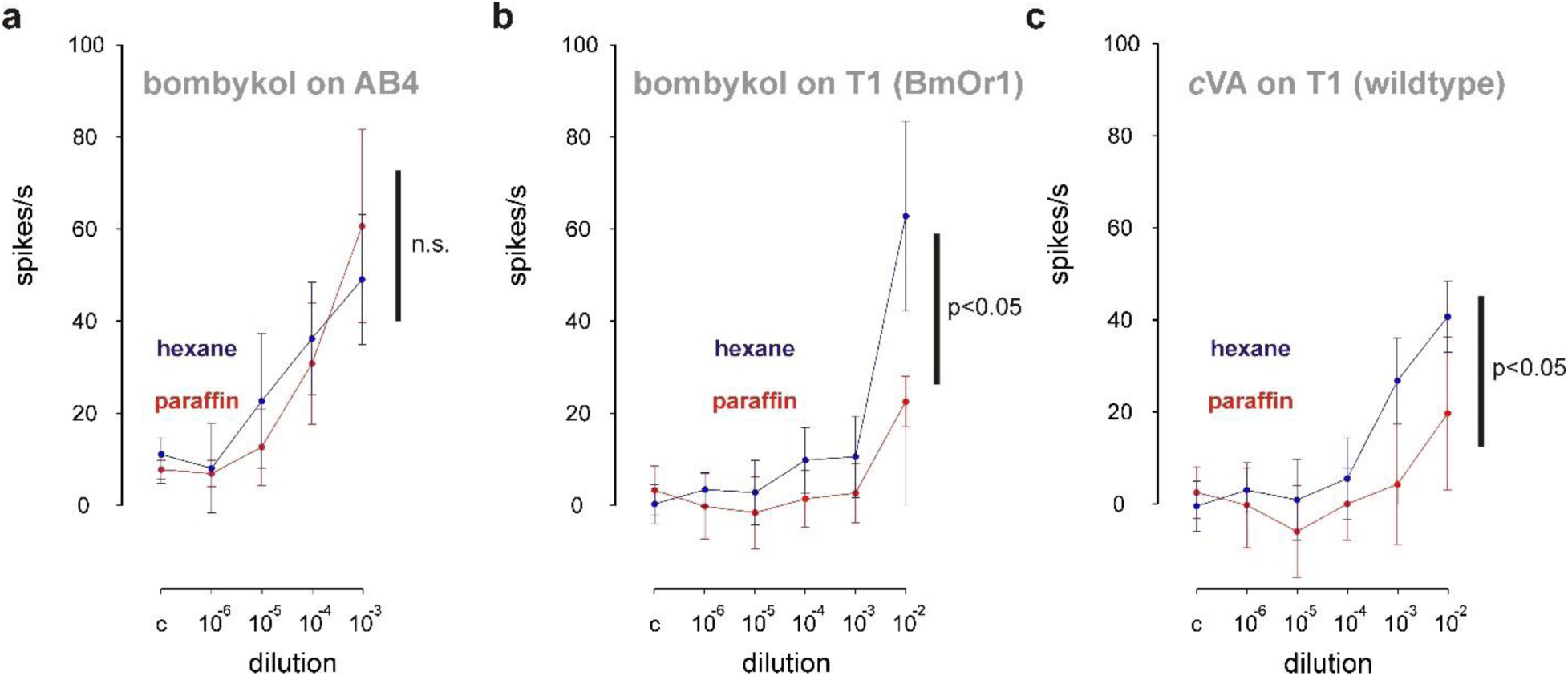
**Using paraffin oil as a solvent instead of hexane suppressed the volatilization of *c*VA and bombykol**, resulting in lower *c*VA-induced responses in Or67d (*c*VA) and bombykol-induced responses in BmOR1-expressing T1 neurons. However, paraffin oil did not significantly suppress responses in AB4a neurons to the bombykol sample. Dose response curves of AB4a (**A**) and T1 (**B,C**) neurons to bombykol (**A,B**) and *c*VA (**C**) diluted in hexane (purple) or paraffin oil (red). Data points represent means (n=7 (**A**) or 8 (**B, C**) different flies) and their 95% confidence intervals. Two-way ANOVAs for Repeated Measurements either indicated significant (p<0.05,) or no significant (n.s.; p>0.05) differences between hexane and paraffin dilutions. Negative response values resulted when AB4a produced fewer spike/s after stimulation compared to pre-puffing control periods (see methods).

**Extended Data Figure 4.**
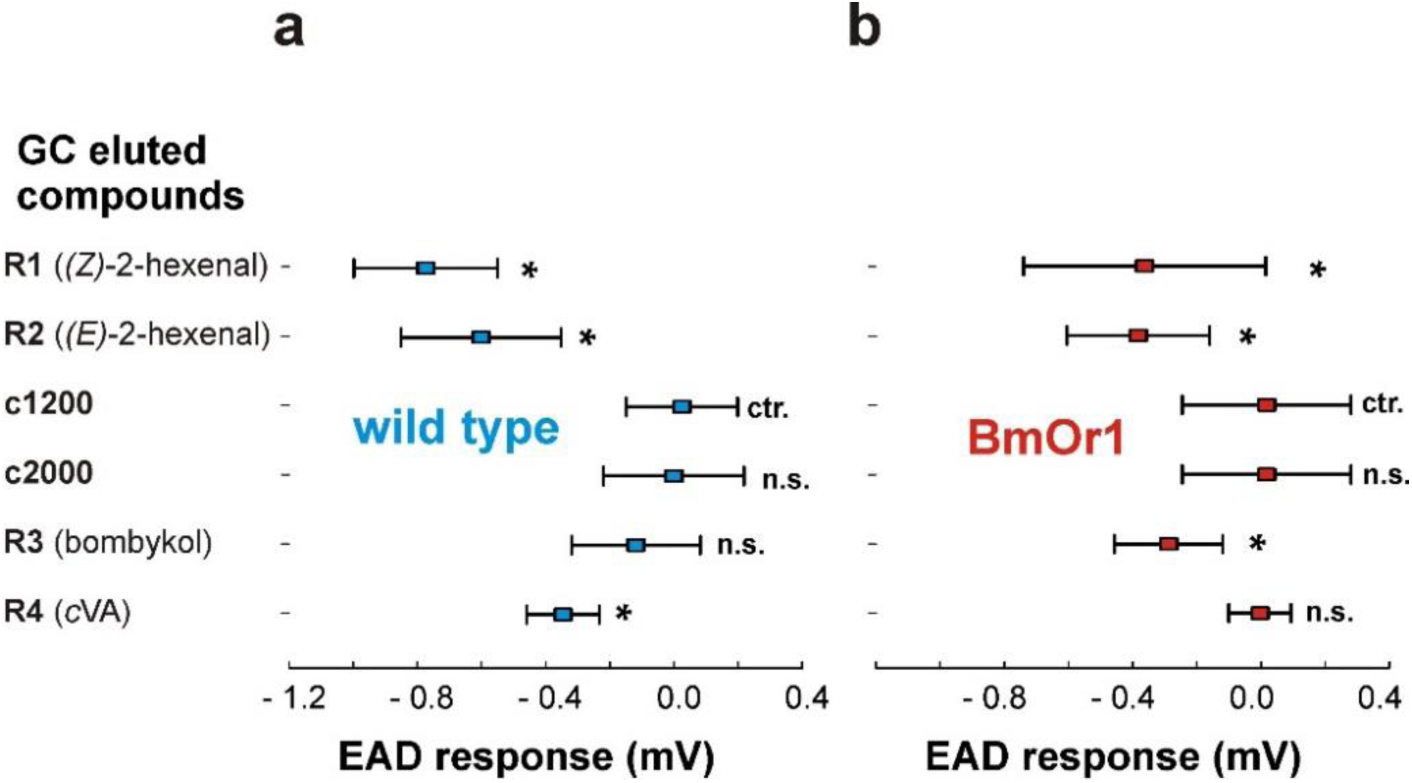
Statistical comparison of *Drosophila* antennal responses (EAD) to compounds eluting off the GC (see Figure 1) relative to controls. **A**, Wildtype fly antennae (n=8 flies), and, **B**, Fly antennae expressing BmOR1 instead of Or67d in T1 neurons (n=8 flies). Responses (means ±95% CI): R1 to (*Z*)-2-hexenal; R2 to (*E*)-2-hexenal; R3 to bombykol; R4 to *c*VA; c1200 and c2000, control responses measured at Kováts Retention Indices 1200 and 2000, respectively. Asterisks (*) indicate significant differences from the control (p<0.05, Holm-Sidak multiple comparisons versus control c1200) following One Way ANOVA analyses (n= 2x 8 different flies). n.s. = not significant.

**Extended Data Figure 5.**
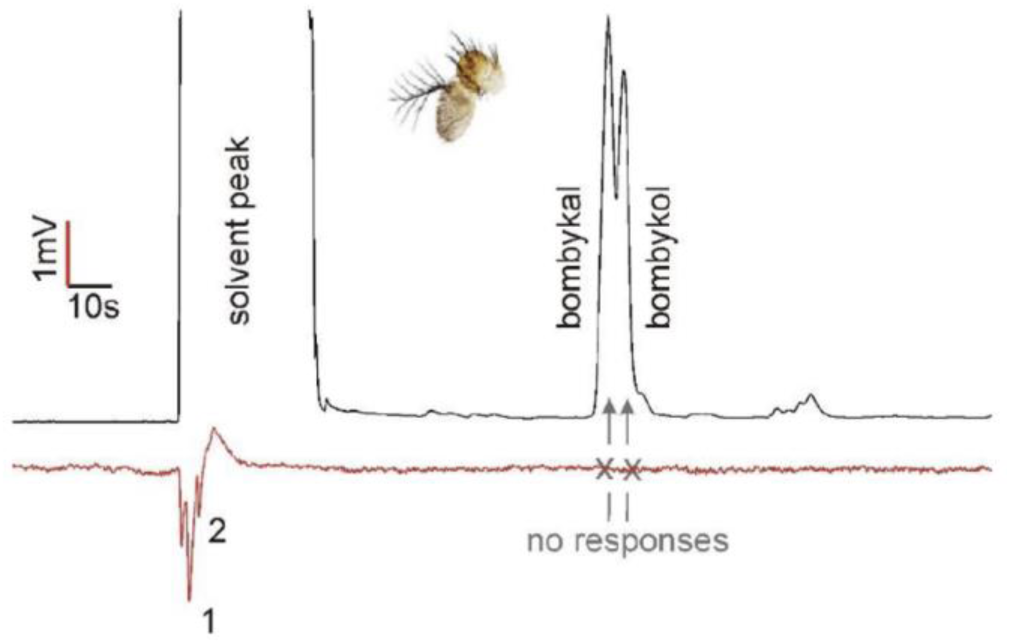
**Responses of wildtype flies in GC-EAD assays** to a blend of bombykol and bombykal (10 µg each in hexane) with a chromatography temperature program adapted for fast screening of low volatility compounds (starting at 260°C, instead of 50°C). Fly antennae consistently responded to early eluting compounds (1,2), but showed no responses (n=7) to either bombykol or bombykal.

**Extended Data Figure 6.**
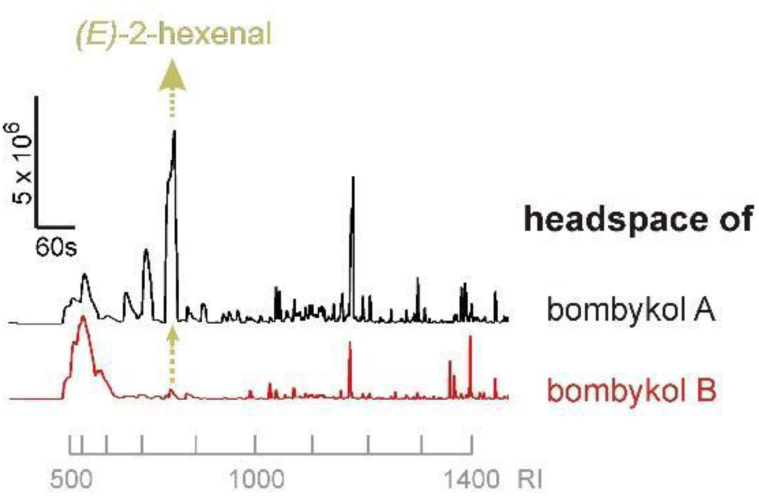
**SPME analysis showing that the headspace of different batches of synthetic bombykol** varied substantially in the quantities of impurities, particularly (*E*)-2-hexenal with the Kováts retention index (RI) of 860.

**Extended Data Figure 7.**
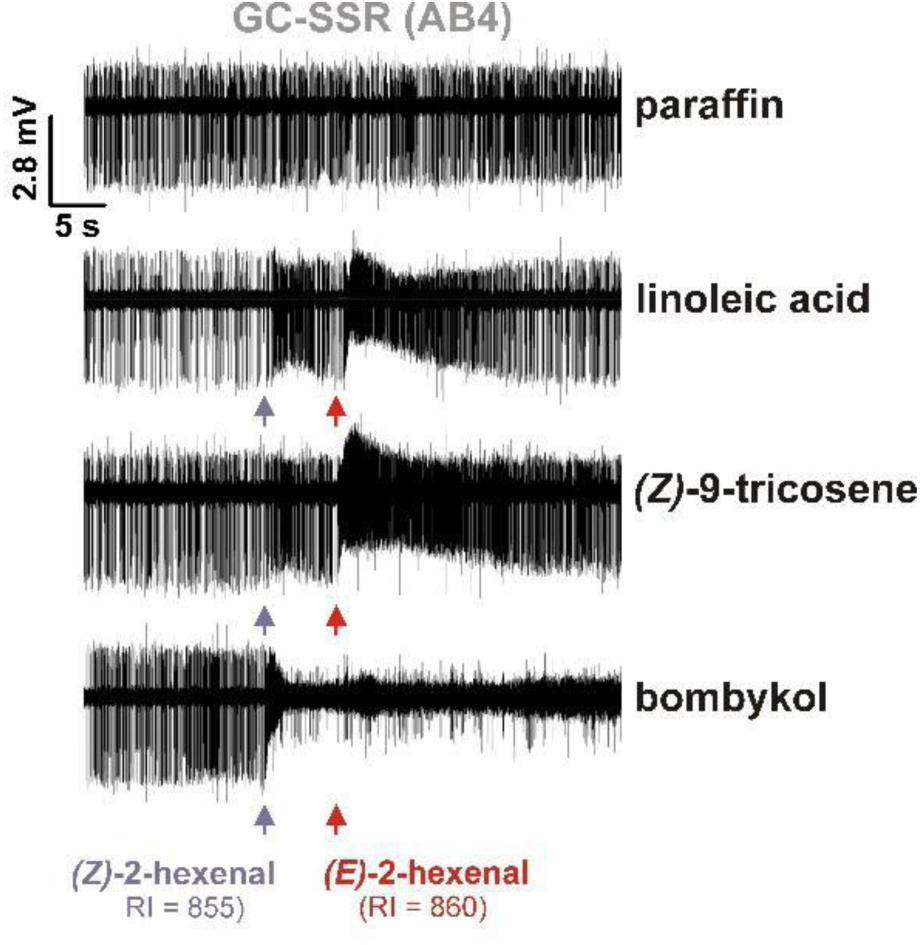
AB4a neurons responded to shared impurities present in the headspace of bombykol, Z9T, and LLA. Aligned segments of GC-SSR traces show that at retention indexes of 885 and 860, injected headspace samples (SPME) of bombykol, Z9T, and LLA contain both isomers of 2-hexenal, which, of all compounds in the injected samples, elicited the largest responses from AB4a neurons.

**Extended Data Figure 8.**
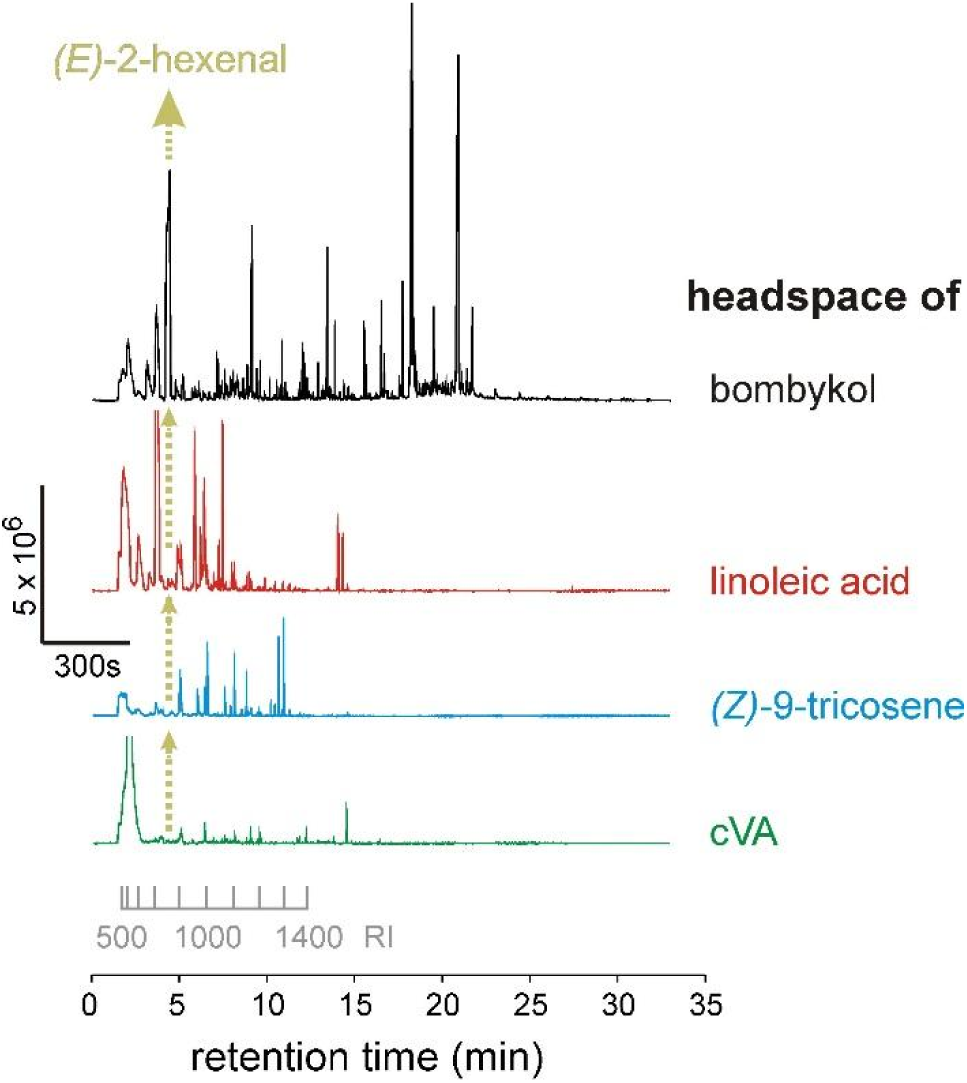
SPME profiles from synthetic standards of bombykol, Z9T, and LLA. The various GC traces demonstrate that the amount of E2H differs substantially between these synthetic standards, and correspondingly the responses of AB4a neurons to these.

**Extended Data Figure 9.**
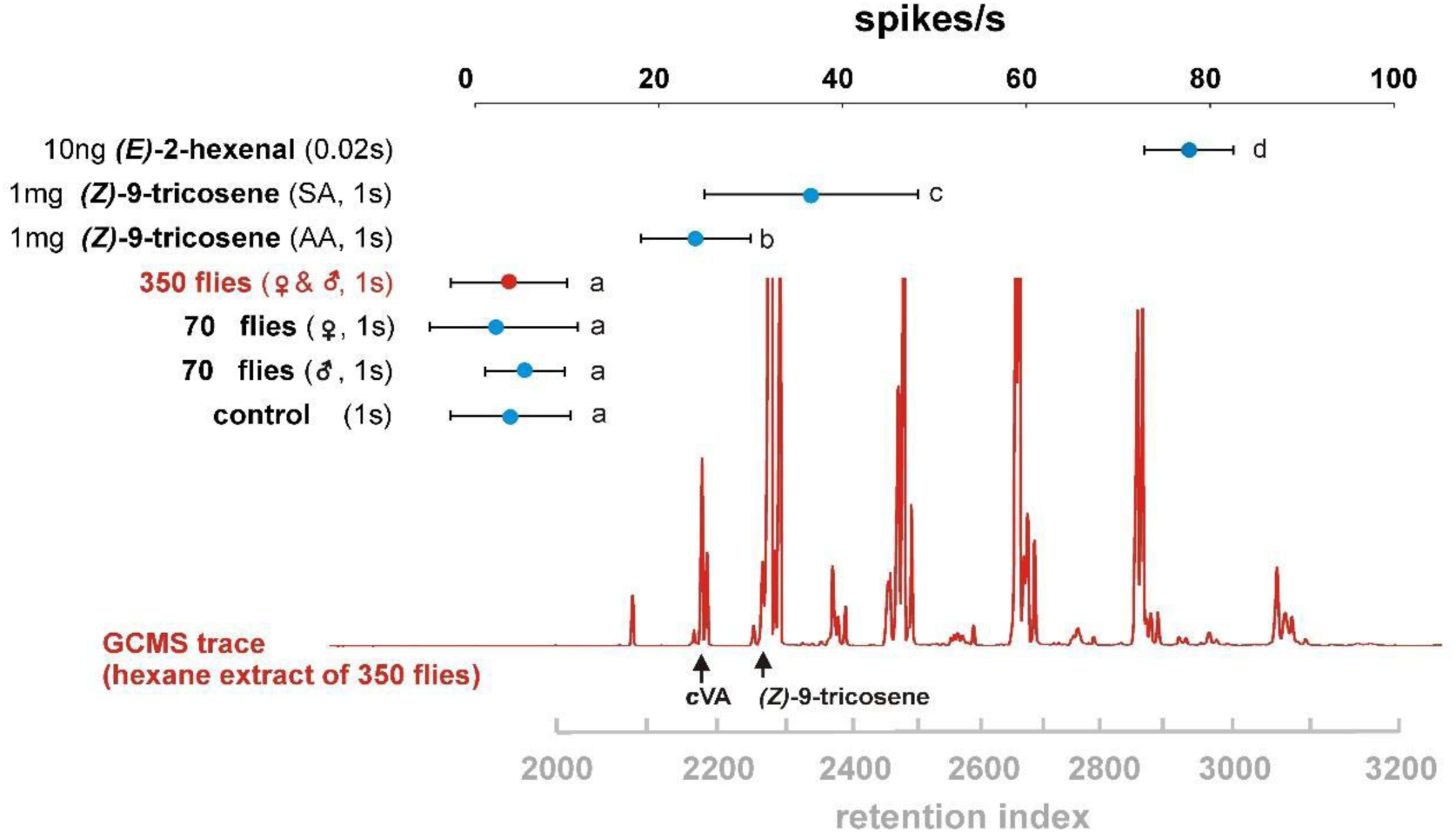
GC-MS profile of cuticular extracts of *Drosophila* and corresponding AB4a neuron responses when puffed with the extracts or with synthetic compounds. Whereas batches of synthetic E2H and Z9T (SIG, Sigma-Aldrich; AA, Alfa Aesar) induced robust responses in AB4a neurons, no responses were observed to a puffed extract of 350 fly equivalents of mixed sex applied on filter paper, to 70 fly equivalents of male or female flies, or to the control (hexane solvent). The solvent was evaporated before the 1s stimulation, E2H lasted for 0.02s. Values (means ±95% confidence intervals) surmounted by the same letters are not significantly different from each other (One Way Repeated Measurements ANOVA, followed by Holm-Sidak multiple comparisons; p>0.05, N=7 different flies). GC-MS trace (red) is from an extract of 350 mixed sex flies.

**Extended Data Figure 10.**
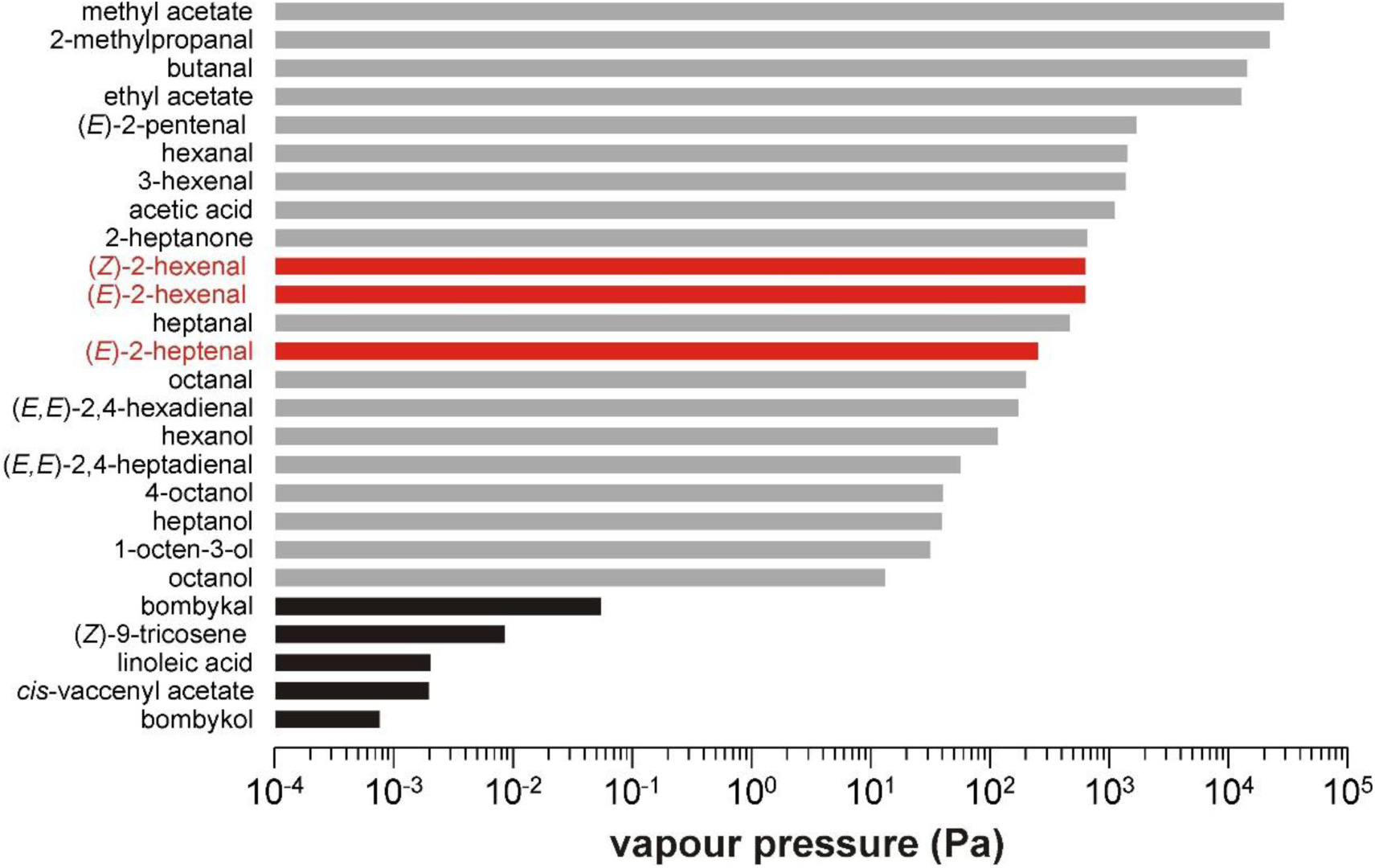
Impurities in synthetic compounds can have dramatically higher vapour pressures than the compounds themselves. The bars represent the calculated values of various impurities (see supplementary table 5) arranged in order of decreasing volatility, and presented on a logarithmic scale. Black bars show the values for the compounds studied in the present work. Grey and red bars show the values for those identified impurities that elicited physiological responses. The three physiologically most important impurities with regard to the antennal basiconic 4a neurons of *D. melanogaster* are shown in red (see also tables 1-3).

## METHODS

### Insects

The wildtype *Drosophila melanogaster* (Meigen 1830; Diptera, Drosophilidae) were originally collected in 2007 from Dalby, Skåne county, Sweden^25^. Transgenic *D. melanogaster* expressing *Bombyx mori* receptor BmOR1 in T1 sensilla were created by crossing OR67d-Gal4 lines (gift from Barry Dickson, Janelia Research Campus, VA, USA) with UAS-BmOR1 (gift from Walter Leal, University of California, Davis CA, USA) to create w;UAS-BmOR1/CyO;OR67d-Gal4/MKRS, which were selected to obtain homozygous flies^26,27,28^ for recordings. Experimental flies were 3-8 days old. Stocks were kept on standard cornmeal-yeast-agar diet under a 12 L:12 D photoperiod and at 22 ^°^C.

*Bombyx mori* (Linnaeus 1758; Lepidoptera, Bombycidae) was obtained either from Agri Pet Garden (Conselve, Italy) or Bugs-World (Budapest, Hungary). Larvae were either fed on a commercial diet (Agri Pet Garden, a mixture of dried & ground mulberry leaves with corn meal, soy meal, agar and water) or on freshly collected mulberry foliage. Larvae were kept in a climatic chamber (26±1 °C, 65±5% RH, 16h light: 8h dark photoperiod) and moved to new boxes every 2-3 days with fresh food. Pupae were removed and placed in boxes lined with paper until eclosion.

### Chemicals and stimuli

Synthetic compounds were acquired from mostly commercial sources with the highest purity available. (10*E*,12*Z*)-10,12-Hexadecadien-1-ol (bombykol, purity >95%), (10*E*,12*Z*)-10,12-hexadecadien-1-al (bombykal, purity >95%), and (*Z*)-11-octadecenyl acetate (*cis-*vaccenyl acetate, *c*VA) were obtained from Pherobank (Wijk bij Duurstede, The Netherlands). (*Z*)-9-Tricosene was purchased from Sigma-Aldrich (Bellefonte, PA, USA) (purity >97%) and Alfa-Aesar (Haverhill, MA, USA) (purity 96%). Other chemicals, including alkane standards for calculation of retention indices (purity ≥99% each), mineral oil (CAS: 8042-47-5), (9*Z*,12*Z*)-9,12-octadecadienoic acid (linoleic acid, LLA, purity 99%) and (*E*)-2-hexenal (98% purity) were obtained from Sigma-Aldrich.

(*Z*)-7-Tricosene was synthesized as follows: sodium hexamethyldisilazide (NaHMDS, 0.75 M in tetrahydrofuran,THF) was added to a slurry of hexadecyltriphenylphosphonium bromide (3.98 g, 7 mmol) in THF at 0ºC under argon until an orange color persisted, followed by addition of another 9.3 ml (7 mmol) of NaHMDS solution. The resulting mixture was stirred 1 h at 0ºC, then cooled to -78ºC in a dry ice-acetone bath, and heptanal (0.74 g, 6.5 mmol) in 4 ml THF was added dropwise. The mixture was allowed to warm to room temp over several hours, then quenched with dilute aqueous NH_4_Cl, and extracted with hexane. The hexane extract was washed with water and brine, dried over anhydrous Na_2_SO_4_, and concentrated. The residue was triturated with hexane, and the soluble portion was purified by vacuum flash chromatography on silica gel, eluting with hexane. The purified product was then recrystallized from acetone at -20ºC overnight, yielding 1.34 g of a white solid which melted at ~0ºC. The recrystallized product contained about 3% of the (*E*)-isomer.

For fly extracts, flies were first freeze-killed (~10 min at -20^°^C) and then placed into a clean 1.5 ml glass vial to which 3 µl of hexane were added per fly (e.g. 1050 µl for 350 flies) at room temperature. The glass was gently shaken for 5 min once a minute and the resulting cuticular extract was transferred to another glass vial. The extract was concentrated by letting the excess hexane evaporate under a gentle nitrogen flow until ~100 µl of the extract remained. The remaining extract was transferred to a 300 µl glass insert, to carefully further concentrate the extract down to 30 µl. The same procedure was followed for the control (hexane without flies).

Stimuli were diluted in either *n*-hexane (Merck, purity ≥99%) or in mineral oil, and 10 µl (30 µl for fly extracts and their controls) of solutions were applied to filter paper disks (12.7 mm Ø; Schleicher 222 & Schnell GmbH, Dassel, Germany) placed inside Pasteur pipettes. Sodium borohydride (Sigma-Aldrich, purity>=96%) was used to reduce the aldehydes present in the batches of 2-hexenal and bombykol to the corresponding alcohols (e.g. (*E*)-2-hexenal to (*E*)-2-hexenol) by adding 25 µl of a saturated solution of NaBH_4_ in ethanol (≥99.9%; Sigma-Aldrich) to 25 µl of hexane solutions of either bombykol (2 mg) or 2-hexenal (4 µg), respectively, at room temperature. After 5 min, the mixtures were diluted with hexane so as to achieve the desired experimental dilutions (bombykol: 5 µg/µl; E2H: 10 ng/µl). We confirmed the successful reduction of the aldehydes by GCMS analysis. Controls consisted of hexane solutions of bombykol and 2-hexenal which were treated only with EtOH. For GCMS assisted verification of (*Z*)-2- hexenal a solution of (*Z*)-2-hexenol (0.1 g, 1 mmol) in 5 ml CH_2_Cl_2_ was cooled in an icebath, and finely powdered pyridinium dichromate (0.56 g, 1.5 mmol) was added in one portion. The mixture was stirred for 2 h at 0ºC, then diluted with 15 ml ether and filtered through a plug of celite filtering aid. The resulting crude product contained an ~1:1 ratio of the (*Z*)- and (*E*)-isomers, as determined by comparison of the retention time of the second isomer with that of an authentic standard or (*E*)-2-hexenal.

### Electrophysiological recordings

For electrophysiological recordings, flies were immobilized in pipette tips with the antennae protruding from the tip. Preparations were placed under a continuous charcoal-filtered, humidified air stream (1 l min^-1^). Stimuli were injected into this airstream using either a stimulus controller (CS-55, Syntech, Kirchzarten, Germany) or as the column effluent from the gas chromatograph (see below). Signals from whole mount antennae (electroantennogram, EAG, using glass electrodes) or single sensilla (SSR, using tungsten electrodes) were collected simultaneously with GC signals (GC-EAD and GC-SSR, respectively), using Syntech hardware and software following established methods^4,28^. Responses of single neurons were expressed as spikes s^-1^ following stimulation, minus the average spike activity in a 1 s prestimulus window. For depletion series, stimulus cartridges were puffed repeatedly into an exhaust vent at 5 s intervals. Other details, such as stimulus load and duration, are noted in the text and figures.

### Headspace collections, gas chromatography (GC), coupled gas chromatography-mass spectrometry (GC-MS) and identifications

Headspace volatiles were collected for 20 min at room temperature using a solid-phase microextraction (SPME, 50/30 μm DVB/CAR/PDMS StableFlex fiber, Supelco/Sigma-Aldrich, Bellefonte, PA) inserted into a 1.5 mL screwcap vial either empty (control) or loaded with 10 µl of the standard being sampled. The fibre was cleaned before use by desorption for 10 min in a 250°C GC injector port. The pre-sampling headspace equilibration time was 10 min. Volatiles collected on the fibre were thermally desorbed directly into the splitless injector of the GC or GC-MS for 0.5 min at 250°C.

GC, GC-EAD, GC-SSR, and GC-MS usually employed a HP-5 capillary column (GC-EAD: 30 m × 0.32 mm × 0.25 µm film; GC-MS: 30 (later 60) m × 0.25 mm × 0.25 µm film, Agilent Technologies, Santa Clara, CA). For Kováts Index-aided verification of structures we also employed DB-WAX columns (either 30 m × 0.32 mm × 0.25 µm film or 60 m × 0.25 mm × 0.25 µm film; Agilent) with helium as carrier gas at 1.84 ml min^-1^. The HP-5 column was used with a temperature program of 50ºC/1 min, then 10ºC min^-1^ to 315ºC, hold for 10 min, whereas the DB-WAX column was programmed from either 35°C/1 min or 30°C/3 min, then 8°C min^-1^ to 230°C for 5 min, unless stated otherwise. For GC-EAD/SSR the effluent was split equally between the flame ionization detector and an antennal preparation, with the portion directed to the antennal preparation exiting via a heated transfer line into a 1 l min^-1^ clean humidified airstream as described above. Electrophysiological and GC signals were integrated using an IDAC-4 A/D converter (Syntech).

Samples were analysed by GC-MS (HP 5890 GC and HP 5975 MS instruments, Agilent) in electron impact ionization mode at 70 eV, scanning a mass range of *m/z* 29–300, at 5 scans s^-1^. Single Ion Monitoring was occasionally employed for trace identification of specific compounds, as specified in the tables. Compounds were tentatively identified by comparison of their mass spectra with mass spectral databases (NIST11 and Wiley) and published Kováts indices, and verified through injection of authentic standards where possible, as specified in the tables.

### Vapour pressure calculations

The vapour pressure values were estimated at 25°C, calculated according to the instructions and methods described in Yaws (2015)^18^. When calculation estimation requirements or missing experimental data did not allow this, estimates were calculated using EPI Suite™ (v4.11, June 2017), a freely available software developed by the EPA’s Office of Pollution Prevention Toxics and Syracuse Research Corporation (SRC Inc.) for the U.S. Environmental Protection Agency.

### Statistical analyses

Statistical analyses described in the text and figure legends were performed two-sided using Sigmastat 4.0 (Systat Software Inc.), or R 3.2.0 software (R Development Core Team 2015). When the assumptions of normality (Shapiro–Wilk test p > 0.05) and equal variance (Spearman rank correlation p > 0.05) were not met, the data (Extended Data Figures 2a, 3b and 3c) were ln –transformed prior to analysis if the assumptions could thus be successfully met.

---

**Extended Data Table 1.**
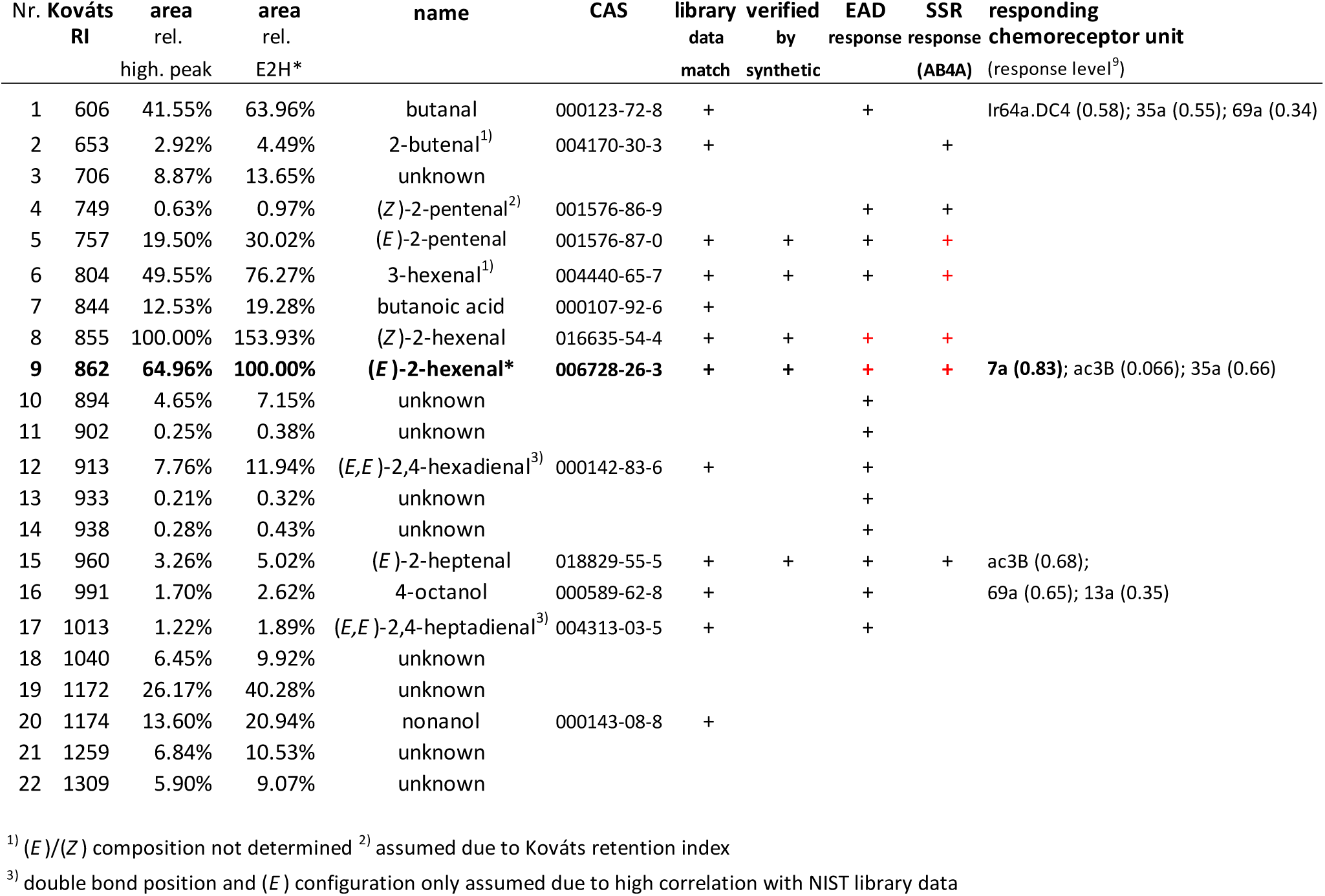
Impurities found in a synthetic standard of bombykol ((10*E*,12*Z*)-10,12-hexadecadienol) by GCMS analysis of headspace volatiles collected by solid phase microextraction. Compounds which were also detected in the control (glass vial without a sample) were excluded if they were not found within treatments in substantially higher amounts (> 200%). Shown are measured Kováts retention indices (RI), percent relative to the most abundant compound, and percent relative to (*E*)-2-hexenal in the sample. Only compounds which either elicited a reproducible electrophysiological response, or occurred in amounts greater than 5% relative to the main compound peak area were included in the table. The asterisk indicates the compound that was mainly responsible for eliciting responses from the *D. melanogaster* AB4a neuron. Compounds were tentatively identified by matches with NIST database spectra, and where possible, identifications were confirmed by matching retention times and mass spectra with those of authentic standards. Positive GC-EAD and GC-SSR responses are indicated; responses in red were also observed with injected samples of bombykol (1 µg). Listed in addition are chemoreceptors reported (DoOR 2.0 database^9^) to respond most strongly to a given compound. The reported response level, ranging from 0 (no response) to 1 (max excitation), is indicated within brackets.

| Nr. | Kováts<br>RI | area<br>rel.<br>high. peak | area<br>rel.<br>E2H* | name | CAS | library<br>data<br>match | verified<br>by<br>synthetic | EAD<br>response | SSR<br>response<br>(AB4A) | responding<br>chemoreceptor unit<br>(response level<sup>9</sup>) |
| --- | --- | --- | --- | --- | --- | --- | --- | --- | --- | --- |
| 1 | 606 | 41.55% | 63.96% | butanal | 000123-72-8 | + |  | + |  | Ir64a.DC4 (0.58); 35a (0.55); 69a (0.34) |
| 2 | 653 | 2.92% | 4.49% | 2-butenal<sup>1)</sup> | 004170-30-3 | + |  |  | + |  |
| 3 | 706 | 8.87% | 13.65% | unknown |  |  |  |  |  |  |
| 4 | 749 | 0.63% | 0.97% | (*Z*)-2-pentenal<sup>2)</sup> | 001576-86-9 |  |  | + | + |  |
| 5 | 757 | 19.50% | 30.02% | (*E*)-2-pentenal | 001576-87-0 | + | + | + | + |  |
| 6 | 804 | 49.55% | 76.27% | 3-hexenal<sup>1)</sup> | 004440-65-7 | + | + | + | + |  |
| 7 | 844 | 12.53% | 19.28% | butanoic acid | 000107-92-6 | + |  |  |  |  |
| 8 | 855 | 100.00% | 153.93% | (*Z*)-2-hexenal | 016635-54-4 | + | + | + | + |  |
| 9 | 862 | 64.96% | 100.00% | (*E*)-2-hexenal* | 006728-26-3 | + | + | + | + | 7a (0.83); ac3B (0.066); 35a (0.66) |
| 10 | 894 | 4.65% | 7.15% | unknown |  |  |  | + |  |  |
| 11 | 902 | 0.25% | 0.38% | unknown |  |  |  | + |  |  |
| 12 | 913 | 7.76% | 11.94% | (*E,E*)-2,4-hexadienal<sup>3)</sup> | 000142-83-6 | + |  | + |  |  |
| 13 | 933 | 0.21% | 0.32% | unknown |  |  |  | + |  |  |
| 14 | 938 | 0.28% | 0.43% | unknown |  |  |  | + |  |  |
| 15 | 960 | 3.26% | 5.02% | (*E*)-2-heptenal | 018829-55-5 | + | + | + | + | ac3B (0.68); |
| 16 | 991 | 1.70% | 2.62% | 4-octanol | 000589-62-8 | + |  | + |  | 69a (0.65); 13a (0.35) |
| 17 | 1013 | 1.22% | 1.89% | (*E,E*)-2,4-heptadienal<sup>3)</sup> | 004313-03-5 | + |  | + |  |  |
| 18 | 1040 | 6.45% | 9.92% | unknown |  |  |  |  |  |  |
| 19 | 1172 | 26.17% | 40.28% | unknown |  |  |  |  |  |  |
| 20 | 1174 | 13.60% | 20.94% | nonanol | 000143-08-8 | + |  |  |  |  |
| 21 | 1259 | 6.84% | 10.53% | unknown |  |  |  |  |  |  |
| 22 | 1309 | 5.90% | 9.07% | unknown |  |  |  |  |  |  |

<sup>1)</sup> (*E*)/(*Z*) composition not determined <sup>2)</sup> assumed due to Kováts retention index
<sup>3)</sup> double bond position and (*E*) configuration only assumed due to high correlation with NIST library data

**Extended Data Table 2.** Impurities found in a synthetic standard of linoleic acid ((9*Z*,12*Z*)-9,12- octadecadienoic acid) by GCMS analysis of headspace volatiles collected by solid phase microextraction. Compounds which were also detected in the control (glass vial without a sample) were excluded if they were not found within treatments in substantially higher amounts (> 200%). Shown are measured Kováts retention indices (RI), percent relative to the most abundant compound, and percent relative to (*E*)-2-hexenal in the bombykol sample (table 1). Only compounds which either elicited a reproducible electrophysiological response, or occurred in amounts greater than 5% relative to the peak area of the main compound were included in the table. The asterisks indicate the compounds that were mainly responsible for eliciting responses from the *D. melanogaster* AB4a neuron. Compounds were tentatively identified by matches with NIST database spectra, and where possible by comparison with authentic standards. Positive GC-EAD and GC-SSR responses are indicated; responses in red were also observed with injected samples of linoleic acid (1 µg). Listed in addition are chemoreceptors reported (DoOR 2.0 database^9^) to respond most strongly to a given compound. The reported response level, ranging from 0 (no response) to 1 (max excitation), is indicated within brackets.

| Nr. | Kováts RI | area rel. high. peak | area rel. E2H* | name | CAS | library data match | verified by synthetic | EAD response | SSR response (AB4A) | responding chemoreceptor unit (response level<sup>9</sup>) |
| --- | --- | --- | --- | --- | --- | --- | --- | --- | --- | --- |
| 1 | 556 | 33.03% | 25.58% | 2-methylpropanal | 000078-84-2 | + |  | + |  |  |
| 2 | 700 | 25.09% | 19.43% | heptane | 000142-82-5 | + |  |  |  |  |
| 3 | 752 | <0.1% | <0.1% | 2-pentenal (traces)<sup>1)</sup> | 000764-39-6 | + |  |  | + |  |
| 4 | 806 | 100.00% | 77.43% | hexanal | 000066-25-1 | + | + | + | + | 85b (0.69); ac3B (0.65); 35a (0.56) |
| 5 | 855 | 0.76% | 0.59% | (*Z*)-2-hexenal | 016635-54-4 | + | + | + | + |  |
| 6 | **856** | **1.94%** | **1.50%** | **(*E*)-2-hexenal\*** | **006728-26-3** | + | + | + | **+** | **7a (0.83);** ac3B (0.066); 35a (0.66) |
| 7 | 870 | 3.15% | 2.44% | hexanol | 000111-27-3 | + |  | + |  |  |
| 8 | 894 | 9.16% | 7.09% | 2-heptanone | 000124-11-8 | + | + | + |  | 85c (0.65); 85b (0.65); 98a (0.62) |
| 9 | 904 | 2.53% | 1.96% | heptanal | 000111-71-7 | + |  | + |  | ac3B (0.59); 22a (0.50); 13a (0.28) |
| 10 | 908 | 7.43% | 5.76% | unknown |  |  |  |  |  |  |
| 11 | 929 | 0.67% | 0.52% | unknown |  |  |  | + |  |  |
| 12 | 948 | 0.52% | 0.41% | (*Z*)-2-heptenal | 057266-86-1 | + |  | + | + |  |
| 13 | **960** | **21.63%** | **16.75%** | **(*E*)-2-heptenal\*** | **018829-55-5** | + | + | + | **+** | ac3B (0.68) |
| 14 | 981 | 8.60% | 6.66% | 1-octen-3-ol | 003391-86-4 | + |  | + |  | 13a (0.79); 85c (0.69); 98a (0.58) |
| 15 | 989 | 1.44% | 1.11% | unknown |  |  |  | + |  |  |
| 16 | 994 | 17.54% | 13.58% | 2-pentyl furan | 003777-69-3 | + |  |  |  |  |
| 17 | 1005 | 2.62% | 2.03% | octanal | 000124-13-0 | + |  | + |  | 35a (0.48); 69a (0.30); ac3B (0.27) |
| 18 | 1062 | 17.56% | 13.60% | 2-octenal<sup>1)</sup> | 002363-89-5 | + |  |  |  |  |

<sup>1)</sup> (*E*)/(*Z*) composition not determined

**Extended Data Table 3.**
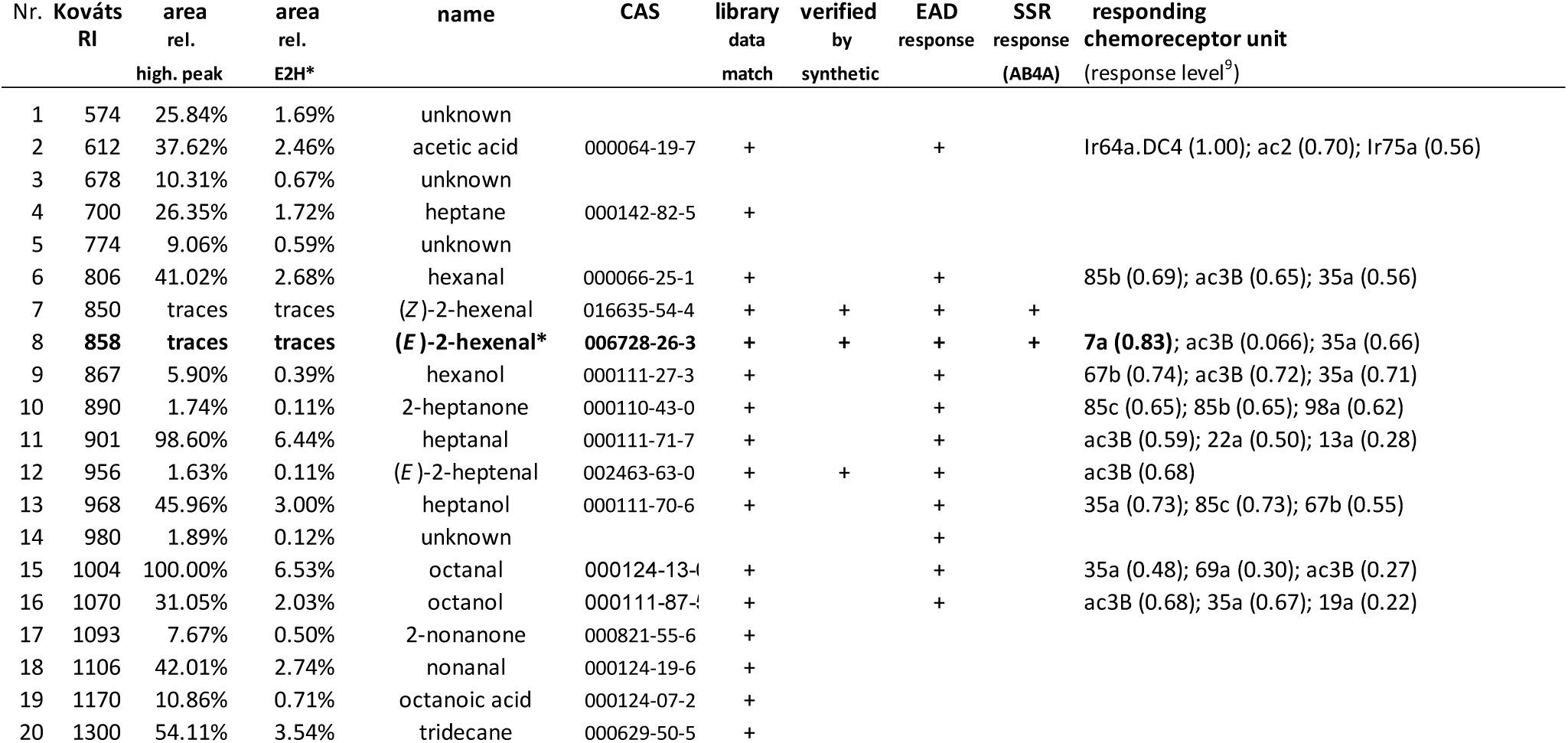
Impurities found in a synthetic standard of (*Z*)-9-tricosene by GCMS analysis of headspace volatiles collected by solid phase microextraction. Compounds which were also detected in the control (glass vial without a sample) were excluded if they were not found within treatments in substantially higher amounts (> 200%). Shown are measured Kováts retention indices (RI), percent relative to the most abundant compound, and percent relative to (*E*)-2-hexenal in the bombykol sample (Table 1). Only compounds which either elicited a reproducible electrophysiological response, or occurred in amounts greater than 5% relative to the peak area of the main compound were included in the table. The asterisk indicates the compound that was mainly responsible for eliciting responses from the *D. melanogaster* AB4a neuron. Compounds were tentatively identified by matches with NIST database spectra, and where possible by comparison with authentic standards. Positive GC-EAD and GC-SSR responses are indicated. Listed in addition are chemoreceptors reported (DoOR 2.0 database^9^) to respond most strongly to a given compound. The reported response level, ranging from 0 (no response) to 1 (max excitation), is indicated within brackets.

**Extended Data Table 4.** Impurities found in a synthetic standard of *cis*-vaccenyl acetate (*c*VA) by GCMS analysis of headspace volatiles collected by solid phase microextraction. Compounds which were also detected in the control (glass vial without a sample) were excluded if they were not found within treatments in substantially higher amounts (> 200%). Shown are measured Kováts retention indices (RI), percent relative to the most abundant compound, and percent relative to (*E*)-2-hexenal in the bombykol sample (Table 1). Only compounds which either elicited a reproducible electrophysiological response, or occurred in amounts greater than 5% relative to the peak area of the main compound were included in the table. The asterisks indicate the volatiles that were mainly responsible for eliciting responses from the *D. melanogaster* antenna. Compounds were tentatively identified by matches with NIST database spectra, and where possible by comparison with authentic standards. Positive GC-EAD responses are indicated. Listed in addition are chemoreceptors reported (DoOR 2.0 database^9^) to respond most strongly to a given compound. The reported response level, ranging from 0 (no response) to 1 (max excitation), is indicated within brackets.

| Nr. | Kováts RI | area rel. high. peak | area rel. E2H* | name | CAS | library data match | verified by synthetic | EAD response | responding chemoreceptor unit (response level<sup>9</sup>) |
| --- | --- | --- | --- | --- | --- | --- | --- | --- | --- |
| 1 | 503 | 10.74% | 6.39% | acetone | 000067-64-1 | + |  |  |  |
| 2 | 542 | traces | traces | methyl acetate | 000079-20-9 | + |  | + | 59b (0.74); 42a (0.61); 42b (0.52) |
| 3 | 600 | 29.35% | 17.47% | hexane | 000110-54-3 | + | + |  |  |
| 4 | 615 | 4.12% | 2.45% | ethyl acetate* | 000141-78-6 | + | + | + | 42a (0.72); 42b (0.61); 59b (0.58) |
| 5 | 618 | 100.00% | 59.53% | acetic acid | 000064-19-7 | + |  | (+)<sup>§</sup> | Ir64a.DC4 (1.00); ac2 (0.70); Ir75a (0.56) |
| 6 | 655 | 10.96% | 6.52% | methyl cyclopentane | 000096-37-7 | + |  |  |  |
| 7 | 676 | 6.02% | 3.59% | cyclohexane | 000110-82-7 | + |  |  |  |
| 8 | 755 | 0.21% | 0.12% | unknown |  |  |  | + |  |
| 9 | 802 | 0.76% | 0.45% | hexanal | 000066-25-1 | + | + | + | 85b (0.69); ac3B (0.65); 35a (0.56) |
| 10 | 850 | 0.58% | 0.34% | (*Z*)-2-hexenal | 016635-54-4 | + | + | + |  |
| 11 | 854 | 1.19% | 0.71% | (*E*)-2-hexenal* | 006728-26-3 | + | + | + | 7a (0.83); ac3B (0.066); 35a (0.66) |

<sup>§</sup>(+) in brackets because acetic acid coelutes together with several other minor compounds on HP5 columns

**Supplementary Table 1.**
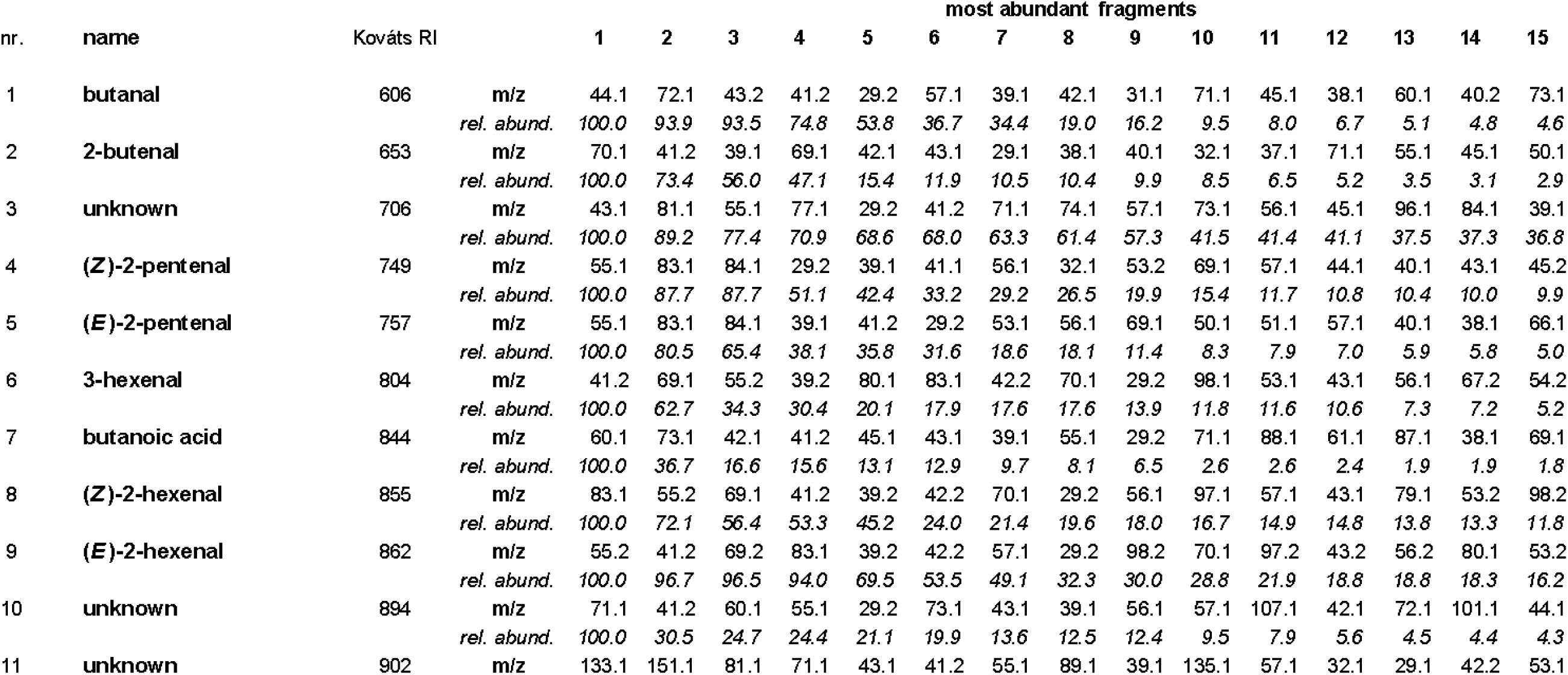
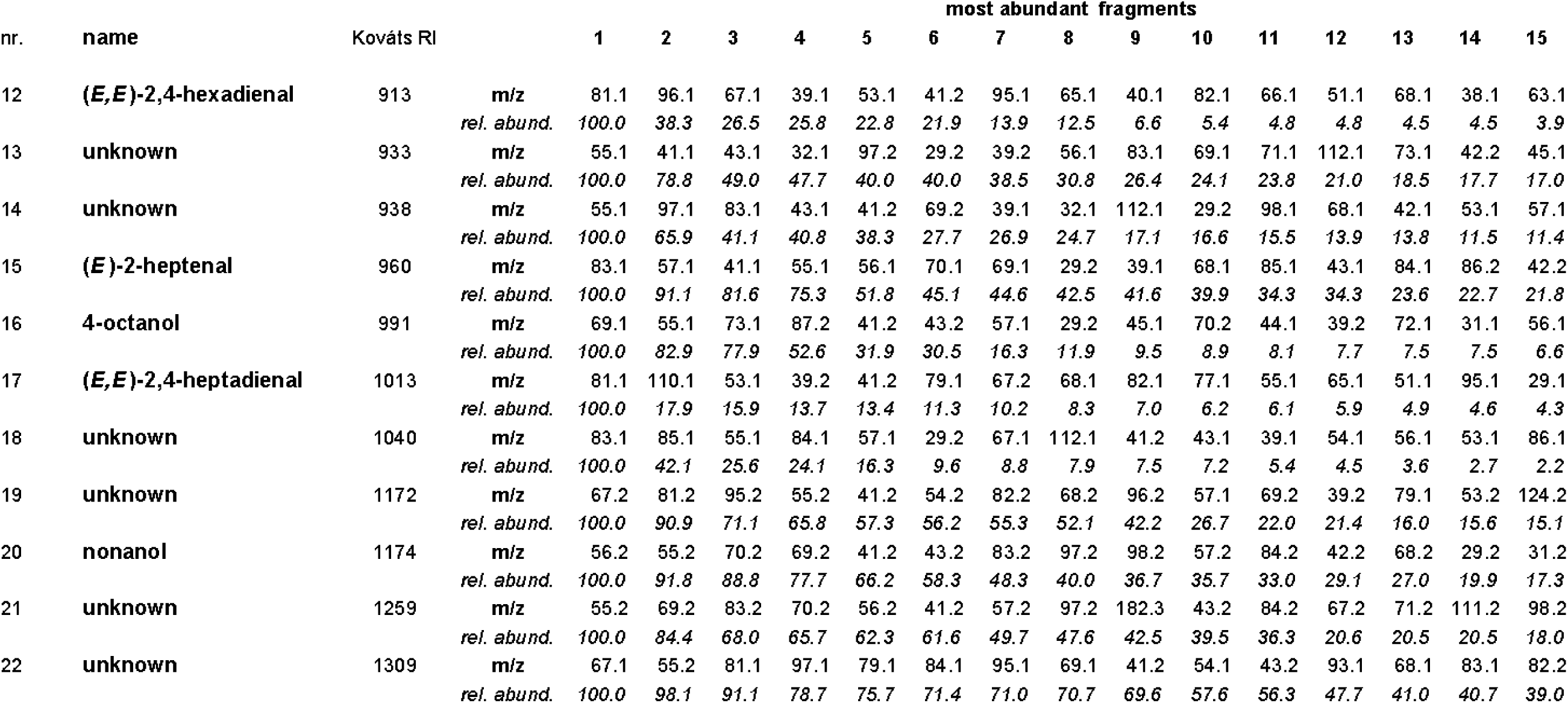
Mass to charge number of bombykol impurities listed in Extended Data Table 1. For each compound, the Kováts retention index (RI) and the fifteen most abundant fragments by their mass to charge numbers are given, followed by each fragment’s abundance relative to the base peak at 100% abundance.

| nr. | name | Kováts RI |  | most abundant fragments |  |  |  |  |  |  |  |  |  |  |  |  |  |  |
| --- | --- | --- | --- | --- | --- | --- | --- | --- | --- | --- | --- | --- | --- | --- | --- | --- | --- | --- |
|  |  |  |  | 1 | 2 | 3 | 4 | 5 | 6 | 7 | 8 | 9 | 10 | 11 | 12 | 13 | 14 | 15 |
| 1 | butanal | 606 | m/z | 44.1 | 72.1 | 43.2 | 41.2 | 29.2 | 57.1 | 39.1 | 42.1 | 31.1 | 71.1 | 45.1 | 38.1 | 60.1 | 40.2 | 73.1 |
|  |  |  | rel. abund. | 100.0 | 93.9 | 93.5 | 74.8 | 53.8 | 36.7 | 34.4 | 19.0 | 16.2 | 9.5 | 8.0 | 6.7 | 5.1 | 4.8 | 4.6 |
| 2 | 2-butenal | 653 | m/z | 70.1 | 41.2 | 39.1 | 69.1 | 42.1 | 43.1 | 29.1 | 38.1 | 40.1 | 32.1 | 37.1 | 71.1 | 55.1 | 45.1 | 50.1 |
|  |  |  | rel. abund. | 100.0 | 73.4 | 56.0 | 47.1 | 15.4 | 11.9 | 10.5 | 10.4 | 9.9 | 8.5 | 6.5 | 5.2 | 3.5 | 3.1 | 2.9 |
| 3 | unknown | 706 | m/z | 43.1 | 81.1 | 55.1 | 77.1 | 29.2 | 41.2 | 71.1 | 74.1 | 57.1 | 73.1 | 56.1 | 45.1 | 96.1 | 84.1 | 39.1 |
|  |  |  | rel. abund. | 100.0 | 89.2 | 77.4 | 70.9 | 68.6 | 68.0 | 63.3 | 61.4 | 57.3 | 41.5 | 41.4 | 41.1 | 37.5 | 37.3 | 36.8 |
| 4 | (Z)-2-pentenal | 749 | m/z | 55.1 | 83.1 | 84.1 | 29.2 | 39.1 | 41.1 | 56.1 | 32.1 | 53.2 | 69.1 | 57.1 | 44.1 | 40.1 | 43.1 | 45.2 |
|  |  |  | rel. abund. | 100.0 | 87.7 | 87.7 | 51.1 | 42.4 | 33.2 | 29.2 | 26.5 | 19.9 | 15.4 | 11.7 | 10.8 | 10.4 | 10.0 | 9.9 |
| 5 | (E)-2-pentenal | 757 | m/z | 55.1 | 83.1 | 84.1 | 39.1 | 41.2 | 29.2 | 53.1 | 56.1 | 69.1 | 50.1 | 51.1 | 57.1 | 40.1 | 38.1 | 66.1 |
|  |  |  | rel. abund. | 100.0 | 80.5 | 65.4 | 38.1 | 35.8 | 31.6 | 18.6 | 18.1 | 11.4 | 8.3 | 7.9 | 7.0 | 5.9 | 5.8 | 5.0 |
| 6 | 3-hexenal | 804 | m/z | 41.2 | 69.1 | 55.2 | 39.2 | 80.1 | 83.1 | 42.2 | 70.1 | 29.2 | 98.1 | 53.1 | 43.1 | 56.1 | 67.2 | 54.2 |
|  |  |  | rel. abund. | 100.0 | 62.7 | 34.3 | 30.4 | 20.1 | 17.9 | 17.6 | 17.6 | 13.9 | 11.8 | 11.6 | 10.6 | 7.3 | 7.2 | 5.2 |
| 7 | butanoic acid | 844 | m/z | 60.1 | 73.1 | 42.1 | 41.2 | 45.1 | 43.1 | 39.1 | 55.1 | 29.2 | 71.1 | 88.1 | 61.1 | 87.1 | 38.1 | 69.1 |
|  |  |  | rel. abund. | 100.0 | 36.7 | 16.6 | 15.6 | 13.1 | 12.9 | 9.7 | 8.1 | 6.5 | 2.6 | 2.6 | 2.4 | 1.9 | 1.9 | 1.8 |
| 8 | (Z)-2-hexenal | 855 | m/z | 83.1 | 55.2 | 69.1 | 41.2 | 39.2 | 42.2 | 70.1 | 29.2 | 56.1 | 97.1 | 57.1 | 43.1 | 79.1 | 53.2 | 98.2 |
|  |  |  | rel. abund. | 100.0 | 72.1 | 56.4 | 53.3 | 45.2 | 24.0 | 21.4 | 19.6 | 18.0 | 16.7 | 14.9 | 14.8 | 13.8 | 13.3 | 11.8 |
| 9 | (E)-2-hexenal | 862 | m/z | 55.2 | 41.2 | 69.2 | 83.1 | 39.2 | 42.2 | 57.1 | 29.2 | 98.2 | 70.1 | 97.2 | 43.2 | 56.2 | 80.1 | 53.2 |
|  |  |  | rel. abund. | 100.0 | 96.7 | 96.5 | 94.0 | 69.5 | 53.5 | 49.1 | 32.3 | 30.0 | 28.8 | 21.9 | 18.8 | 18.8 | 18.3 | 16.2 |
| 10 | unknown | 894 | m/z | 71.1 | 41.2 | 60.1 | 55.1 | 29.2 | 73.1 | 43.1 | 39.1 | 56.1 | 57.1 | 107.1 | 42.1 | 72.1 | 101.1 | 44.1 |
|  |  |  | rel. abund. | 100.0 | 30.5 | 24.7 | 24.4 | 21.1 | 19.9 | 13.6 | 12.5 | 12.4 | 9.5 | 7.9 | 5.6 | 4.5 | 4.4 | 4.3 |
| 11 | unknown | 902 | m/z | 133.1 | 151.1 | 81.1 | 71.1 | 43.1 | 41.2 | 55.1 | 89.1 | 39.1 | 135.1 | 57.1 | 32.1 | 29.1 | 42.2 | 53.1 |
|  |  |  |  | 1 | 2 | 3 | 4 | 5 | 6 | 7 | 8 | 9 | 10 | 11 | 12 | 13 | 14 | 15 |
| 12 | **(E,E)-2,4-hexadienal** | 913 | **m/z** | 81.1 | 96.1 | 67.1 | 39.1 | 53.1 | 41.2 | 95.1 | 65.1 | 40.1 | 82.1 | 66.1 | 51.1 | 68.1 | 38.1 | 63.1 |
|  |  |  | **rel. abund.** | 100.0 | 38.3 | 26.5 | 25.8 | 22.8 | 21.9 | 13.9 | 12.5 | 6.6 | 5.4 | 4.8 | 4.8 | 4.5 | 4.5 | 3.9 |
| 13 | **unknown** | 933 | **m/z** | 55.1 | 41.1 | 43.1 | 32.1 | 97.2 | 29.2 | 39.2 | 56.1 | 83.1 | 69.1 | 71.1 | 112.1 | 73.1 | 42.2 | 45.1 |
|  |  |  | **rel. abund.** | 100.0 | 78.8 | 49.0 | 47.7 | 40.0 | 40.0 | 38.5 | 30.8 | 26.4 | 24.1 | 23.8 | 21.0 | 18.5 | 17.7 | 17.0 |
| 14 | **unknown** | 938 | **m/z** | 55.1 | 97.1 | 83.1 | 43.1 | 41.2 | 69.2 | 39.1 | 32.1 | 112.1 | 29.2 | 98.1 | 68.1 | 42.1 | 53.1 | 57.1 |
|  |  |  | **rel. abund.** | 100.0 | 65.9 | 41.1 | 40.8 | 38.3 | 27.7 | 26.9 | 24.7 | 17.1 | 16.6 | 15.5 | 13.9 | 13.8 | 11.5 | 11.4 |
| 15 | **(E)-2-heptenal** | 960 | **m/z** | 83.1 | 57.1 | 41.1 | 55.1 | 56.1 | 70.1 | 69.1 | 29.2 | 39.1 | 68.1 | 85.1 | 43.1 | 84.1 | 86.2 | 42.2 |
|  |  |  | **rel. abund.** | 100.0 | 91.1 | 81.6 | 75.3 | 51.8 | 45.1 | 44.6 | 42.5 | 41.6 | 39.9 | 34.3 | 34.3 | 23.6 | 22.7 | 21.8 |
| 16 | **4-octanol** | 991 | **m/z** | 69.1 | 55.1 | 73.1 | 87.2 | 41.2 | 43.2 | 57.1 | 29.2 | 45.1 | 70.2 | 44.1 | 39.2 | 72.1 | 31.1 | 56.1 |
|  |  |  | **rel. abund.** | 100.0 | 82.9 | 77.9 | 52.6 | 31.9 | 30.5 | 16.3 | 11.9 | 9.5 | 8.9 | 8.1 | 7.7 | 7.5 | 7.5 | 6.6 |
| 17 | **(E,E)-2,4-heptadienal** | 1013 | **m/z** | 81.1 | 110.1 | 53.1 | 39.2 | 41.2 | 79.1 | 67.2 | 68.1 | 82.1 | 77.1 | 55.1 | 65.1 | 51.1 | 95.1 | 29.1 |
|  |  |  | **rel. abund.** | 100.0 | 17.9 | 15.9 | 13.7 | 13.4 | 11.3 | 10.2 | 8.3 | 7.0 | 6.2 | 6.1 | 5.9 | 4.9 | 4.6 | 4.3 |
| 18 | **unknown** | 1040 | **m/z** | 83.1 | 85.1 | 55.1 | 84.1 | 57.1 | 29.2 | 67.1 | 112.1 | 41.2 | 43.1 | 39.1 | 54.1 | 56.1 | 53.1 | 86.1 |
|  |  |  | **rel. abund.** | 100.0 | 42.1 | 25.6 | 24.1 | 16.3 | 9.6 | 8.8 | 7.9 | 7.5 | 7.2 | 5.4 | 4.5 | 3.6 | 2.7 | 2.2 |
| 19 | **unknown** | 1172 | **m/z** | 67.2 | 81.2 | 95.2 | 55.2 | 41.2 | 54.2 | 82.2 | 68.2 | 96.2 | 57.1 | 69.2 | 39.2 | 79.1 | 53.2 | 124.2 |
|  |  |  | **rel. abund.** | 100.0 | 90.9 | 71.1 | 65.8 | 57.3 | 56.2 | 55.3 | 52.1 | 42.2 | 26.7 | 22.0 | 21.4 | 16.0 | 15.6 | 15.1 |
| 20 | **nonanol** | 1174 | **m/z** | 56.2 | 55.2 | 70.2 | 69.2 | 41.2 | 43.2 | 83.2 | 97.2 | 98.2 | 57.2 | 84.2 | 42.2 | 68.2 | 29.2 | 31.2 |
|  |  |  | **rel. abund.** | 100.0 | 91.8 | 88.8 | 77.7 | 66.2 | 58.3 | 48.3 | 40.0 | 36.7 | 35.7 | 33.0 | 29.1 | 27.0 | 19.9 | 17.3 |
| 21 | **unknown** | 1259 | **m/z** | 55.2 | 69.2 | 83.2 | 70.2 | 56.2 | 41.2 | 57.2 | 97.2 | 182.3 | 43.2 | 84.2 | 67.2 | 71.2 | 111.2 | 98.2 |
|  |  |  | **rel. abund.** | 100.0 | 84.4 | 68.0 | 65.7 | 62.3 | 61.6 | 49.7 | 47.6 | 42.5 | 39.5 | 36.3 | 20.6 | 20.5 | 20.5 | 18.0 |
| 22 | **unknown** | 1309 | **m/z** | 67.1 | 55.2 | 81.1 | 97.1 | 79.1 | 84.1 | 95.1 | 69.1 | 41.2 | 54.1 | 43.2 | 93.1 | 68.1 | 83.1 | 82.2 |
|  |  |  | **rel. abund.** | 100.0 | 98.1 | 91.1 | 78.7 | 75.7 | 71.4 | 71.0 | 70.7 | 69.6 | 57.6 | 56.3 | 47.7 | 41.0 | 40.7 | 39.0 |

**Supplementary Table 2.**
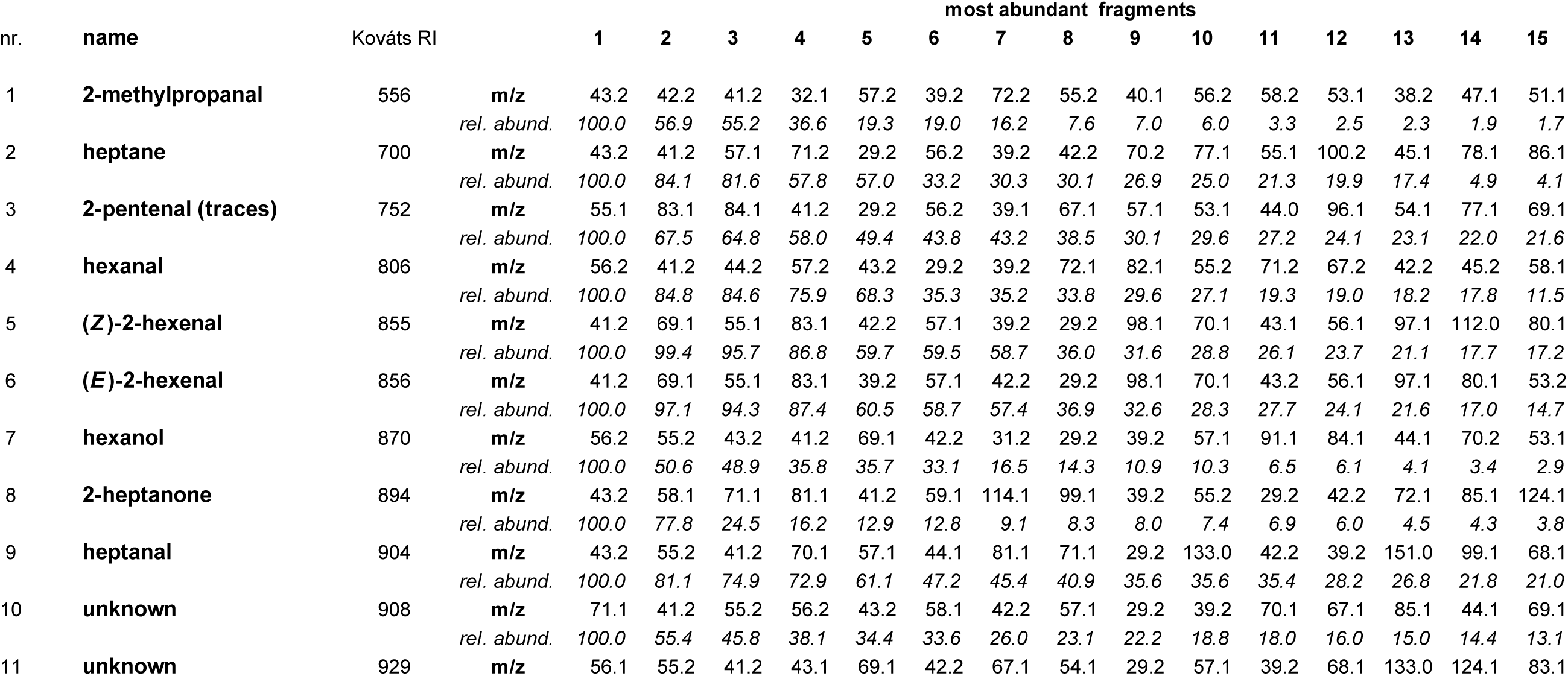
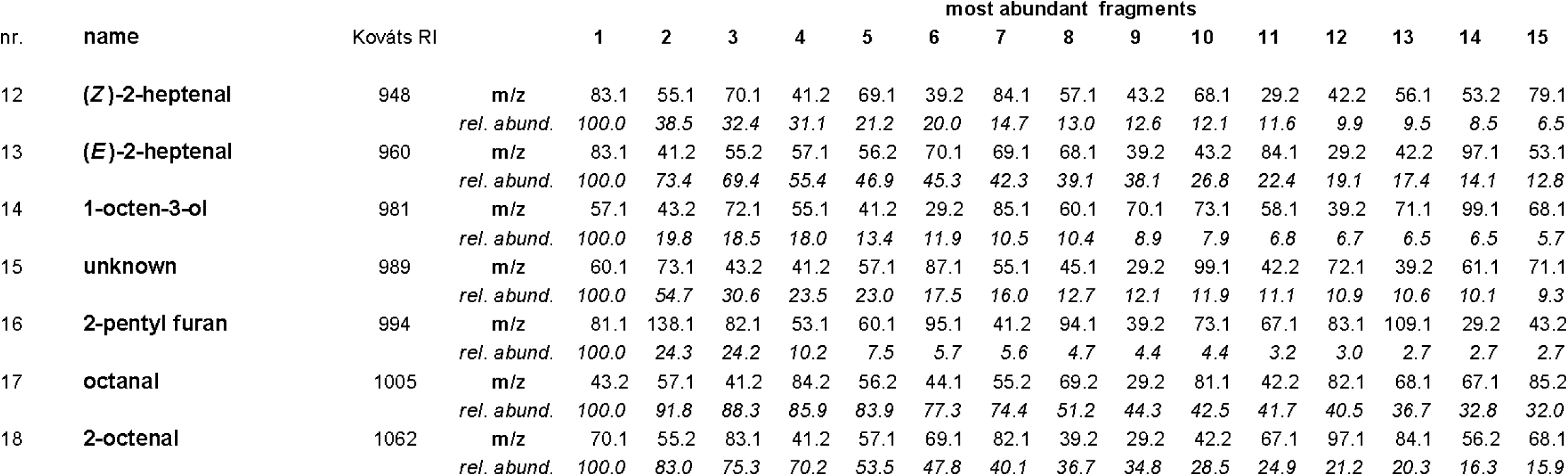
Mass to charge ratios of ions from linoleic acid impurities listed in Extended Data Table 2. For each compound, the Kováts retention index (RI) and the fifteen most abundant fragments by their mass to charge numbers are given, followed by each fragment’s abundance relative to the base peak at 100% abundance.

| nr. | name | Kováts RI |  | most abundant fragments |  |  |  |  |  |  |  |  |  |  |  |  |  |  |
| --- | --- | --- | --- | --- | --- | --- | --- | --- | --- | --- | --- | --- | --- | --- | --- | --- | --- | --- |
|  |  |  |  | 1 | 2 | 3 | 4 | 5 | 6 | 7 | 8 | 9 | 10 | 11 | 12 | 13 | 14 | 15 |
| 1 | 2-methylpropanal | 556 | m/z | 43.2 | 42.2 | 41.2 | 32.1 | 57.2 | 39.2 | 72.2 | 55.2 | 40.1 | 56.2 | 58.2 | 53.1 | 38.2 | 47.1 | 51.1 |
|  |  |  | rel. abund. | 100.0 | 56.9 | 55.2 | 36.6 | 19.3 | 19.0 | 16.2 | 7.6 | 7.0 | 6.0 | 3.3 | 2.5 | 2.3 | 1.9 | 1.7 |
| 2 | heptane | 700 | m/z | 43.2 | 41.2 | 57.1 | 71.2 | 29.2 | 56.2 | 39.2 | 42.2 | 70.2 | 77.1 | 55.1 | 100.2 | 45.1 | 78.1 | 86.1 |
|  |  |  | rel. abund. | 100.0 | 84.1 | 81.6 | 57.8 | 57.0 | 33.2 | 30.3 | 30.1 | 26.9 | 25.0 | 21.3 | 19.9 | 17.4 | 4.9 | 4.1 |
| 3 | 2-pentenal (traces) | 752 | m/z | 55.1 | 83.1 | 84.1 | 41.2 | 29.2 | 56.2 | 39.1 | 67.1 | 57.1 | 53.1 | 44.0 | 96.1 | 54.1 | 77.1 | 69.1 |
|  |  |  | rel. abund. | 100.0 | 67.5 | 64.8 | 58.0 | 49.4 | 43.8 | 43.2 | 38.5 | 30.1 | 29.6 | 27.2 | 24.1 | 23.1 | 22.0 | 21.6 |
| 4 | hexanal | 806 | m/z | 56.2 | 41.2 | 44.2 | 57.2 | 43.2 | 29.2 | 39.2 | 72.1 | 82.1 | 55.2 | 71.2 | 67.2 | 42.2 | 45.2 | 58.1 |
|  |  |  | rel. abund. | 100.0 | 84.8 | 84.6 | 75.9 | 68.3 | 35.3 | 35.2 | 33.8 | 29.6 | 27.1 | 19.3 | 19.0 | 18.2 | 17.8 | 11.5 |
| 5 | (Z)-2-hexenal | 855 | m/z | 41.2 | 69.1 | 55.1 | 83.1 | 42.2 | 57.1 | 39.2 | 29.2 | 98.1 | 70.1 | 43.1 | 56.1 | 97.1 | 112.0 | 80.1 |
|  |  |  | rel. abund. | 100.0 | 99.4 | 95.7 | 86.8 | 59.7 | 59.5 | 58.7 | 36.0 | 31.6 | 28.8 | 26.1 | 23.7 | 21.1 | 17.7 | 17.2 |
| 6 | (E)-2-hexenal | 856 | m/z | 41.2 | 69.1 | 55.1 | 83.1 | 39.2 | 57.1 | 42.2 | 29.2 | 98.1 | 70.1 | 43.2 | 56.1 | 97.1 | 80.1 | 53.2 |
|  |  |  | rel. abund. | 100.0 | 97.1 | 94.3 | 87.4 | 60.5 | 58.7 | 57.4 | 36.9 | 32.6 | 28.3 | 27.7 | 24.1 | 21.6 | 17.0 | 14.7 |
| 7 | hexanol | 870 | m/z | 56.2 | 55.2 | 43.2 | 41.2 | 69.1 | 42.2 | 31.2 | 29.2 | 39.2 | 57.1 | 91.1 | 84.1 | 44.1 | 70.2 | 53.1 |
|  |  |  | rel. abund. | 100.0 | 50.6 | 48.9 | 35.8 | 35.7 | 33.1 | 16.5 | 14.3 | 10.9 | 10.3 | 6.5 | 6.1 | 4.1 | 3.4 | 2.9 |
| 8 | 2-heptanone | 894 | m/z | 43.2 | 58.1 | 71.1 | 81.1 | 41.2 | 59.1 | 114.1 | 99.1 | 39.2 | 55.2 | 29.2 | 42.2 | 72.1 | 85.1 | 124.1 |
|  |  |  | rel. abund. | 100.0 | 77.8 | 24.5 | 16.2 | 12.9 | 12.8 | 9.1 | 8.3 | 8.0 | 7.4 | 6.9 | 6.0 | 4.5 | 4.3 | 3.8 |
| 9 | heptanal | 904 | m/z | 43.2 | 55.2 | 41.2 | 70.1 | 57.1 | 44.1 | 81.1 | 71.1 | 29.2 | 133.0 | 42.2 | 39.2 | 151.0 | 99.1 | 68.1 |
|  |  |  | rel. abund. | 100.0 | 81.1 | 74.9 | 72.9 | 61.1 | 47.2 | 45.4 | 40.9 | 35.6 | 35.6 | 35.4 | 28.2 | 26.8 | 21.8 | 21.0 |
| 10 | unknown | 908 | m/z | 71.1 | 41.2 | 55.2 | 56.2 | 43.2 | 58.1 | 42.2 | 57.1 | 29.2 | 39.2 | 70.1 | 67.1 | 85.1 | 44.1 | 69.1 |
|  |  |  | rel. abund. | 100.0 | 55.4 | 45.8 | 38.1 | 34.4 | 33.6 | 26.0 | 23.1 | 22.2 | 18.8 | 18.0 | 16.0 | 15.0 | 14.4 | 13.1 |
| 11 | unknown | 929 | m/z | 56.1 | 55.2 | 41.2 | 43.1 | 69.1 | 42.2 | 67.1 | 54.1 | 29.2 | 57.1 | 39.2 | 68.1 | 133.0 | 124.1 | 83.1 |
|  |  |  |  | 1 | 2 | 3 | 4 | 5 | 6 | 7 | 8 | 9 | 10 | 11 | 12 | 13 | 14 | 15 |
| 12 | (Z)-2-heptenal | 948 | m/z | 83.1 | 55.1 | 70.1 | 41.2 | 69.1 | 39.2 | 84.1 | 57.1 | 43.2 | 68.1 | 29.2 | 42.2 | 56.1 | 53.2 | 79.1 |
|  |  |  | rel. abund. | 100.0 | 38.5 | 32.4 | 31.1 | 21.2 | 20.0 | 14.7 | 13.0 | 12.6 | 12.1 | 11.6 | 9.9 | 9.5 | 8.5 | 6.5 |
| 13 | (E)-2-heptenal | 960 | m/z | 83.1 | 41.2 | 55.2 | 57.1 | 56.2 | 70.1 | 69.1 | 68.1 | 39.2 | 43.2 | 84.1 | 29.2 | 42.2 | 97.1 | 53.1 |
|  |  |  | rel. abund. | 100.0 | 73.4 | 69.4 | 55.4 | 46.9 | 45.3 | 42.3 | 39.1 | 38.1 | 26.8 | 22.4 | 19.1 | 17.4 | 14.1 | 12.8 |
| 14 | 1-octen-3-ol | 981 | m/z | 57.1 | 43.2 | 72.1 | 55.1 | 41.2 | 29.2 | 85.1 | 60.1 | 70.1 | 73.1 | 58.1 | 39.2 | 71.1 | 99.1 | 68.1 |
|  |  |  | rel. abund. | 100.0 | 19.8 | 18.5 | 18.0 | 13.4 | 11.9 | 10.5 | 10.4 | 8.9 | 7.9 | 6.8 | 6.7 | 6.5 | 6.5 | 5.7 |
| 15 | unknown | 989 | m/z | 60.1 | 73.1 | 43.2 | 41.2 | 57.1 | 87.1 | 55.1 | 45.1 | 29.2 | 99.1 | 42.2 | 72.1 | 39.2 | 61.1 | 71.1 |
|  |  |  | rel. abund. | 100.0 | 54.7 | 30.6 | 23.5 | 23.0 | 17.5 | 16.0 | 12.7 | 12.1 | 11.9 | 11.1 | 10.9 | 10.6 | 10.1 | 9.3 |
| 16 | 2-pentyl furan | 994 | m/z | 81.1 | 138.1 | 82.1 | 53.1 | 60.1 | 95.1 | 41.2 | 94.1 | 39.2 | 73.1 | 67.1 | 83.1 | 109.1 | 29.2 | 43.2 |
|  |  |  | rel. abund. | 100.0 | 24.3 | 24.2 | 10.2 | 7.5 | 5.7 | 5.6 | 4.7 | 4.4 | 4.4 | 3.2 | 3.0 | 2.7 | 2.7 | 2.7 |
| 17 | octanal | 1005 | m/z | 43.2 | 57.1 | 41.2 | 84.2 | 56.2 | 44.1 | 55.2 | 69.2 | 29.2 | 81.1 | 42.2 | 82.1 | 68.1 | 67.1 | 85.2 |
|  |  |  | rel. abund. | 100.0 | 91.8 | 88.3 | 85.9 | 83.9 | 77.3 | 74.4 | 51.2 | 44.3 | 42.5 | 41.7 | 40.5 | 36.7 | 32.8 | 32.0 |
| 18 | 2-octenal | 1062 | m/z | 70.1 | 55.2 | 83.1 | 41.2 | 57.1 | 69.1 | 82.1 | 39.2 | 29.2 | 42.2 | 67.1 | 97.1 | 84.1 | 56.2 | 68.1 |
|  |  |  | rel. abund. | 100.0 | 83.0 | 75.3 | 70.2 | 53.5 | 47.8 | 40.1 | 36.7 | 34.8 | 28.5 | 24.9 | 21.2 | 20.3 | 16.3 | 15.9 |

**Supplementary Table 3.** Mass to charge ratios of ions from (*Z*)-9-tricosene impurities listed in Extended Data Table 3. For each compound, the Kováts retention index (RI) and the fifteen most abundant fragments by their mass to charge numbers are given, followed by each fragment’s abundance relative to the base peak at 100% abundance.

| nr. | name | Kováts RI |  | most abundant fragments |  |  |  |  |  |  |  |  |  |  |  |  |  |  |
| --- | --- | --- | --- | --- | --- | --- | --- | --- | --- | --- | --- | --- | --- | --- | --- | --- | --- | --- |
|  |  |  |  | 1 | 2 | 3 | 4 | 5 | 6 | 7 | 8 | 9 | 10 | 11 | 12 | 13 | 14 | 15 |
| 1 | unknown | 574 | m/z | 44.1 | 40.1 | 43.1 | 29.2 | 45.1 | 60.1 | 59.2 | 42.1 | 41.2 | 58.2 | 75.1 | 207.0 | 34.1 | 39.2 | 57.2 |
|  |  |  | rel. abund. | 100.0 | 98.1 | 57.4 | 45.4 | 42.2 | 30.6 | 30.4 | 9.8 | 8.7 | 7.8 | 6.6 | 6.1 | 4.9 | 4.2 | 4.2 |
| 2 | acetic acid | 612 | m/z | 43.2 | 32.1 | 45.1 | 57.2 | 60.1 | 41.2 | 29.2 | 42.2 | 59.1 | 56.2 | 44.1 | 39.2 | 75.1 | 31.2 | 86.2 |
|  |  |  | rel. abund. | 100.0 | 58.6 | 56.9 | 40.6 | 40.2 | 37.9 | 27.3 | 26.9 | 23.9 | 21.1 | 18.1 | 12.2 | 10.1 | 8.7 | 8.0 |
| 3 | unknown | 678 | m/z | 77.1 | 45.1 | 43.2 | 56.2 | 84.1 | 78.1 | 41.2 | 74.1 | 207.0 | 55.1 | 42.2 | 29.2 | 79.1 | 57.1 | 73.1 |
|  |  |  | rel. abund. | 100.0 | 16.8 | 13.4 | 10.8 | 7.9 | 7.8 | 7.5 | 5.5 | 5.0 | 4.8 | 4.5 | 4.5 | 4.4 | 3.5 | 3.4 |
| 4 | heptane | 700 | m/z | 77.1 | 45.1 | 43.2 | 56.2 | 84.1 | 78.1 | 41.2 | 74.1 | 207.0 | 55.1 | 42.2 | 29.2 | 79.1 | 57.1 | 73.1 |
|  |  |  | rel. abund. | 100.0 | 16.8 | 13.4 | 10.8 | 7.9 | 7.8 | 7.5 | 5.5 | 5.0 | 4.8 | 4.5 | 4.5 | 4.4 | 3.5 | 3.4 |
| 5 | unknown | 774 | m/z | 43.2 | 41.2 | 57.1 | 44.2 | 56.2 | 71.2 | 29.2 | 45.1 | 55.2 | 70.2 | 42.2 | 58.1 | 39.2 | 100.2 | 207.0 |
|  |  |  | rel. abund. | 100.0 | 80.5 | 70.9 | 69.5 | 57.7 | 53.1 | 51.7 | 36.1 | 36.1 | 34.9 | 34.5 | 32.2 | 31.4 | 16.5 | 16.1 |
| 6 | hexanal | 806 | m/z | 91.1 | 60.1 | 92.1 | 41.2 | 55.2 | 42.2 | 70.2 | 73.1 | 43.1 | 39.2 | 29.2 | 45.2 | 69.1 | 57.1 | 31.2 |
|  |  |  | rel. abund. | 100.0 | 92.8 | 58.3 | 51.7 | 51.4 | 50.4 | 37.5 | 32.0 | 31.3 | 23.4 | 21.5 | 21.0 | 18.3 | 17.3 | 15.0 |
| 7 | (Z)-2-hexenal<sup>§</sup> | 850 | m/z | 69.1 | 83.1 | 41.2 | 55.1 | 97.1 | 29.2 | 98.1 |  |  |  |  |  |  |  |  |
|  |  |  | rel. abund. | 100.0 | 93.2 | 78.1 | 76.6 | 44.4 | 25.8 | 6.0 |  |  |  |  |  |  |  |  |
| 8 | (E)-2-hexenal<sup>§</sup> | 858 | m/z | 41.2 | 55.1 | 69.1 | 83.1 | 29.2 | 98.1 | 97.1 |  |  |  |  |  |  |  |  |
|  |  |  | rel. abund. | 100.0 | 82.0 | 70.3 | 49.3 | 44.2 | 34.4 | 24.7 |  |  |  |  |  |  |  |  |
| 9 | hexanol | 867 | m/z | 56.2 | 55.2 | 43.2 | 41.2 | 69.1 | 42.2 | 31.2 | 29.2 | 39.2 | 57.2 | 84.1 | 71.1 | 70.2 | 44.1 | 91.1 |
|  |  |  | rel. abund. | 100.0 | 56.2 | 49.6 | 38.7 | 35.9 | 31.1 | 16.4 | 13.2 | 12.0 | 10.9 | 7.6 | 5.8 | 4.9 | 4.8 | 3.9 |
| 10 | 2-heptanone | 890 | m/z | 43.2 | 58.1 | 56.2 | 41.2 | 55.2 | 70.1 | 71.1 | 39.2 | 42.1 | 29.2 | 97.1 | 114.1 | 98.1 | 44.1 | 54.2 |
|  |  |  | rel. abund. | 100.0 | 55.6 | 39.8 | 39.2 | 36.0 | 21.9 | 20.1 | 18.6 | 16.0 | 15.1 | 12.3 | 7.7 | 6.6 | 4.7 | 4.6 |

<sup>§</sup> Ion fragment values by SIM (Selected Ion Monitoring)

| nr. | name | Kováts RI |  | most abundant fragments |  |  |  |  |  |  |  |  |  |  |  |  |  |  |
| --- | --- | --- | --- | --- | --- | --- | --- | --- | --- | --- | --- | --- | --- | --- | --- | --- | --- | --- |
|  |  |  |  | 1 | 2 | 3 | 4 | 5 | 6 | 7 | 8 | 9 | 10 | 11 | 12 | 13 | 14 | 15 |
| 11 | heptanal | 901 | m/z | 70.2 | 44.2 | 43.2 | 41.2 | 55.2 | 57.1 | 42.2 | 71.1 | 29.2 | 81.1 | 39.2 | 86.1 | 68.1 | 96.1 | 45.2 |
|  |  |  | rel. abund. | 100.0 | 72.4 | 63.5 | 58.6 | 56.8 | 50.3 | 39.3 | 30.8 | 30.5 | 28.7 | 22.1 | 22.0 | 19.5 | 16.3 | 15.9 |
| 12 | (E)-2-heptenal | 956 | m/z | 56.1 | 83.1 | 41.2 | 55.1 | 57.1 | 43.1 | 69.1 | 68.1 | 70.2 | 39.1 | 29.2 | 85.1 | 42.2 | 91.1 | 32.1 |
|  |  |  | rel. abund. | 100.0 | 99.1 | 98.0 | 91.5 | 73.6 | 70.0 | 52.1 | 50.0 | 48.5 | 46.5 | 41.4 | 34.7 | 31.2 | 23.7 | 19.8 |
| 13 | heptanol | 968 | m/z | 70.2 | 56.2 | 55.2 | 69.2 | 41.2 | 43.2 | 42.2 | 57.2 | 29.2 | 31.2 | 68.2 | 39.2 | 83.2 | 71.1 | 54.2 |
|  |  |  | rel. abund. | 100.0 | 78.9 | 62.0 | 52.5 | 50.0 | 45.5 | 32.1 | 20.3 | 15.5 | 15.4 | 14.8 | 11.9 | 9.5 | 6.1 | 4.9 |
| 14 | unknown | 980 | m/z | 83.1 | 41.1 | 55.1 | 39.2 | 43.2 | 133.0 | 103.1 | 94.1 | 59.1 | 60.1 | 87.1 | 70.1 | 45.1 | 75.1 | 113.1 |
|  |  |  | rel. abund. | 100.0 | 48.7 | 47.0 | 31.9 | 24.6 | 23.1 | 21.5 | 21.1 | 19.9 | 19.1 | 19.0 | 18.8 | 18.6 | 17.8 | 17.4 |
| 15 | octanal | 1004 | m/z | 43.2 | 57.2 | 84.2 | 41.2 | 56.2 | 44.1 | 55.2 | 69.1 | 29.2 | 81.1 | 82.1 | 42.2 | 68.1 | 67.1 | 85.2 |
|  |  |  | rel. abund. | 100.0 | 94.6 | 92.1 | 89.2 | 88.7 | 79.7 | 74.1 | 51.6 | 44.8 | 42.7 | 42.5 | 42.2 | 39.7 | 34.5 | 32.4 |
| 16 | octanol | 1070 | m/z | 56.2 | 55.2 | 70.1 | 69.2 | 41.2 | 84.2 | 43.2 | 83.2 | 42.2 | 57.2 | 60.1 | 68.1 | 29.2 | 73.1 | 39.2 |
|  |  |  | rel. abund. | 100.0 | 90.8 | 75.5 | 74.7 | 72.0 | 59.0 | 57.8 | 53.8 | 42.6 | 37.9 | 28.6 | 23.5 | 23.4 | 20.7 | 18.3 |
| 17 | 2-nonanone | 1093 | m/z | 58.1 | 43.1 | 71.1 | 57.2 | 59.2 | 41.2 | 142.2 | 55.2 | 85.1 | 29.2 | 39.2 | 84.1 | 42.2 | 82.1 | 56.2 |
|  |  |  | rel. abund. | 100.0 | 71.1 | 25.1 | 22.1 | 22.0 | 15.6 | 10.4 | 9.0 | 6.3 | 6.1 | 5.8 | 5.3 | 5.0 | 4.9 | 4.3 |
| 18 | nonanal | 1106 | m/z | 57.2 | 41.2 | 56.2 | 43.2 | 55.2 | 98.2 | 44.1 | 70.2 | 82.1 | 69.2 | 68.2 | 81.1 | 95.1 | 29.2 | 96.1 |
|  |  |  | rel. abund. | 100.0 | 62.1 | 58.9 | 51.8 | 49.4 | 43.4 | 43.1 | 41.6 | 37.1 | 35.7 | 32.7 | 29.5 | 29.0 | 28.5 | 27.5 |
| 19 | octanoic acid | 1170 | m/z | 60.1 | 73.1 | 43.2 | 101.1 | 55.2 | 41.2 | 85.2 | 84.1 | 87.1 | 69.2 | 115.1 | 61.1 | 29.2 | 57.2 | 45.1 |
|  |  |  | rel. abund. | 100.0 | 78.1 | 35.1 | 31.2 | 27.7 | 27.3 | 22.0 | 21.2 | 15.7 | 13.8 | 13.1 | 12.6 | 11.9 | 10.1 | 9.9 |
| 20 | tridecane | 1300 | m/z | 57.2 | 71.2 | 43.2 | 85.2 | 41.2 | 55.2 | 56.2 | 70.2 | 29.2 | 99.2 | 184.2 | 84.2 | 42.2 | 69.2 | 98.2 |
|  |  |  | rel. abund. | 100.0 | 69.6 | 62.1 | 47.1 | 28.0 | 15.5 | 15.4 | 13.8 | 10.6 | 9.4 | 8.8 | 8.8 | 8.2 | 7.6 | 7.5 |

**Supplementary Table 4.** Mass to charge ratios of ions from *c*VA impurities listed in Extended Data Table 4. For each compound, the Kováts retention index (RI) and the fifteen most abundant fragments by their mass to charge numbers are given, followed by each fragment’s abundance relative to the base peak at 100% abundance.

| nr. | name | Kováts RI |  | most abundant fragments |  |  |  |  |  |  |  |  |  |  |  |  |  |  |
| --- | --- | --- | --- | --- | --- | --- | --- | --- | --- | --- | --- | --- | --- | --- | --- | --- | --- | --- |
|  |  |  |  | 1 | 2 | 3 | 4 | 5 | 6 | 7 | 8 | 9 | 10 | 11 | 12 | 13 | 14 | 15 |
| 1 | methyl acetate | 542 | m/z | 43.1 | 74.1 | 32.1 | 59.1 | 42.1 | 29.1 | 44.1 | 31.1 | 84.0 | 45.1 | 41.1 | 40.0 | 75.0 | 86.0 | 49.1 |
|  |  |  | rel. abund. | 100.0 | 28.0 | 12.0 | 9.5 | 9.4 | 4.6 | 3.6 | 2.1 | 1.9 | 1.7 | 1.6 | 1.5 | 1.2 | 1.2 | 0.9 |
| 2 | hexane | 600 | m/z | 57.2 | 43.2 | 41.2 | 29.2 | 27.1 | 56.2 | 42.2 | 39.2 | 86.1 | 28.1 | 55.2 | 15.0 | 71.2 | 58.1 | 26.1 |
|  |  |  | rel.abund. | 100.0 | 81.3 | 70.2 | 60.7 | 45.4 | 45.3 | 40.9 | 19.7 | 15.5 | 10.7 | 6.6 | 5.8 | 5.0 | 4.4 | 3.9 |
| 3 | ethyl acetate | 615 | m/z | 43.1 | 45.1 | 60.1 | 15.1 | 42.1 | 29.1 | 14.1 | 28.2 | 41.1 | 18.1 | 44.1 | 31.2 | 16.1 | 13.1 | 61.1 |
|  |  |  | rel.abund. | 100.0 | 90.4 | 74.8 | 17.0 | 13.0 | 8.4 | 4.8 | 4.0 | 3.5 | 2.7 | 2.4 | 2.4 | 2.3 | 2.0 | 1.9 |
| 4 | acetic acid | 618 | m/z | 43.1 | 45.1 | 60.1 | 15.1 | 42.1 | 14.1 | 29.1 | 28.2 | 13.1 | 16.1 | 18.1 | 44.1 | 41.1 | 31.1 | 17.1 |
|  |  |  | rel.abund. | 100.0 | 87.5 | 57.1 | 41.6 | 13.7 | 13.3 | 13.2 | 7.3 | 6.4 | 6.4 | 5.6 | 4.7 | 4.4 | 3.6 | 2.8 |
| 5 | methyl cyclopentane | 655 | m/z | 56.2 | 41.2 | 69.1 | 42.2 | 55.2 | 84.2 | 39.2 | 43.2 | 27.1 | 29.2 | 57.2 | 68.1 | 40.2 | 54.2 | 53.1 |
|  |  |  | rel.abund. | 100.0 | 49.4 | 35.3 | 24.6 | 23.0 | 17.4 | 13.9 | 11.1 | 9.9 | 7.1 | 6.0 | 4.0 | 3.5 | 3.1 | 2.7 |
| 6 | cyclohexane | 676 | m/z | 56.1 | 84.1 | 41.2 | 55.2 | 69.1 | 42.1 | 43.1 | 39.2 | 57.2 | 27.1 | 85.1 | 29.2 | 28.1 | 83.1 | 54.2 |
|  |  |  | rel.abund. | 100.0 | 71.4 | 50.5 | 30.9 | 25.8 | 25.0 | 23.0 | 14.9 | 14.0 | 11.6 | 8.5 | 8.4 | 6.1 | 4.8 | 4.8 |
| 7 | unknown | 755 | m/z | 44.1 | 55.1 | 83.1 | 45.2 | 41.2 | 71.1 | 75.1 | 84.1 | 29.2 | 39.2 | 43.2 | 32.1 | 46.1 | 77.1 | 69.1 |
|  |  |  | rel.abund. | 100.0 | 74.3 | 55.9 | 55.6 | 54.9 | 51.4 | 50.0 | 43.4 | 43.3 | 37.6 | 33.3 | 28.4 | 26.9 | 26.6 | 22.3 |
| 8 | hexanal | 802 | m/z | 44.1 | 56.2 | 41.2 | 43.1 | 57.1 | 27.1 | 29.2 | 39.1 | 45.1 | 72.1 | 55.1 | 82.1 | 42.1 | 58.1 | 67.1 |
|  |  |  | rel.abund. | 100.0 | 82.0 | 69.1 | 55.1 | 38.0 | 33.8 | 32.9 | 20.0 | 19.4 | 16.6 | 15.2 | 12.7 | 10.7 | 8.9 | 8.0 |
| 9 | (Z)-2-hexenal | 850 | m/z | 83.1 | 55.2 | 69.1 | 41.2 | 39.2 | 42.2 | 70.1 | 29.2 | 56.1 | 97.1 | 57.1 | 43.1 | 79.1 | 53.2 | 98.2 |
|  |  |  | rel.abund. | 100.0 | 72.1 | 56.4 | 53.3 | 45.2 | 24.0 | 21.4 | 19.6 | 18.0 | 16.7 | 14.9 | 14.8 | 13.8 | 13.3 | 11.8 |
| 10 | (E)-2-hexenal | 854 | m/z | 41.2 | 55.1 | 42.2 | 69.1 | 39.2 | 83.1 | 57.1 | 27.1 | 29.2 | 98.1 | 43.1 | 70.2 | 56.2 | 53.2 | 80.1 |
|  |  |  | rel.abund. | 100.0 | 61.1 | 58.1 | 57.1 | 50.1 | 48.0 | 47.0 | 43.0 | 42.0 | 26.0 | 23.0 | 17.0 | 14.0 | 9.8 | 9.6 |

**Supplementary Table 5.**
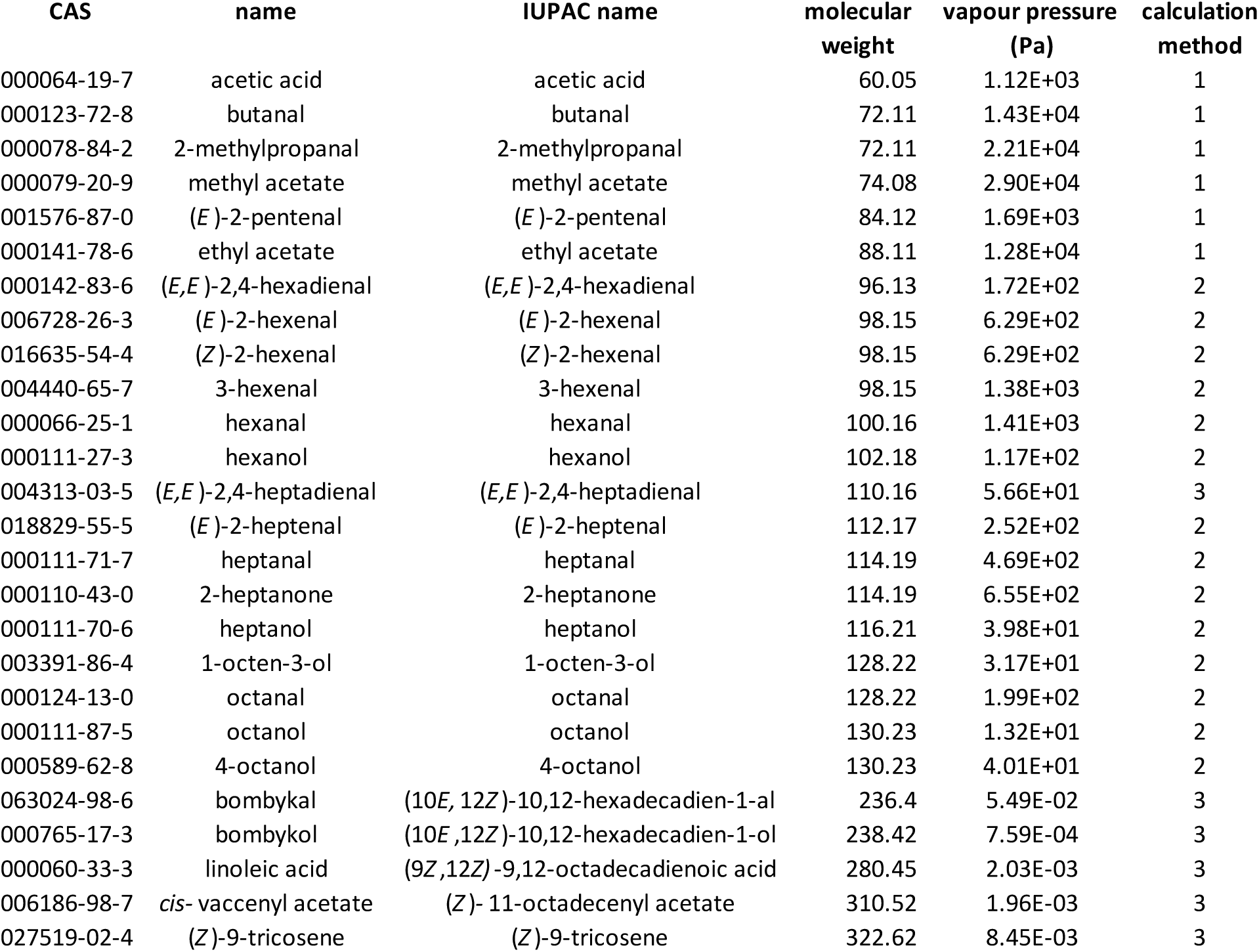
**Calculated vapour pressure** in Pascals for selected compounds of particular importance in the present study (see tables 1-3), arranged by their molecular weights. The vapour pressure values are calculated estimates based either on experimental data provided in literature (method 1) or estimates calculated with the EPI Suite™ software (U.S. Environmental Protection Agency, v4.11, June 2017; method 2 = mean values of Antoine and Grain methods; method 3 = values by the modified Grain method).

| CAS | name | IUPAC name | molecular weight | vapour pressure (Pa) | calculation method |
| --- | --- | --- | --- | --- | --- |
| 000064-19-7 | acetic acid | acetic acid | 60.05 | 1.12E+03 | 1 |
| 000123-72-8 | butanal | butanal | 72.11 | 1.43E+04 | 1 |
| 000078-84-2 | 2-methylpropanal | 2-methylpropanal | 72.11 | 2.21E+04 | 1 |
| 000079-20-9 | methyl acetate | methyl acetate | 74.08 | 2.90E+04 | 1 |
| 001576-87-0 | (E)-2-pentenal | (E)-2-pentenal | 84.12 | 1.69E+03 | 1 |
| 000141-78-6 | ethyl acetate | ethyl acetate | 88.11 | 1.28E+04 | 1 |
| 000142-83-6 | (E,E)-2,4-hexadienal | (E,E)-2,4-hexadienal | 96.13 | 1.72E+02 | 2 |
| 006728-26-3 | (E)-2-hexenal | (E)-2-hexenal | 98.15 | 6.29E+02 | 2 |
| 016635-54-4 | (Z)-2-hexenal | (Z)-2-hexenal | 98.15 | 6.29E+02 | 2 |
| 004440-65-7 | 3-hexenal | 3-hexenal | 98.15 | 1.38E+03 | 2 |
| 000066-25-1 | hexanal | hexanal | 100.16 | 1.41E+03 | 2 |
| 000111-27-3 | hexanol | hexanol | 102.18 | 1.17E+02 | 2 |
| 004313-03-5 | (E,E)-2,4-heptadienal | (E,E)-2,4-heptadienal | 110.16 | 5.66E+01 | 3 |
| 018829-55-5 | (E)-2-heptenal | (E)-2-heptenal | 112.17 | 2.52E+02 | 2 |
| 000111-71-7 | heptanal | heptanal | 114.19 | 4.69E+02 | 2 |
| 000110-43-0 | 2-heptanone | 2-heptanone | 114.19 | 6.55E+02 | 2 |
| 000111-70-6 | heptanol | heptanol | 116.21 | 3.98E+01 | 2 |
| 003391-86-4 | 1-octen-3-ol | 1-octen-3-ol | 128.22 | 3.17E+01 | 2 |
| 000124-13-0 | octanal | octanal | 128.22 | 1.99E+02 | 2 |
| 000111-87-5 | octanol | octanol | 130.23 | 1.32E+01 | 2 |
| 000589-62-8 | 4-octanol | 4-octanol | 130.23 | 4.01E+01 | 2 |
| 063024-98-6 | bombykal | (10E, 12Z)-10,12-hexadecadien-1-al | 236.4 | 5.49E-02 | 3 |
| 000765-17-3 | bombykol | (10E, 12Z)-10,12-hexadecadien-1-ol | 238.42 | 7.59E-04 | 3 |
| 000060-33-3 | linoleic acid | (9Z, 12Z)-9,12-octadecadienoic acid | 280.45 | 2.03E-03 | 3 |
| 006186-98-7 | cis- vaccenyl acetate | (Z)- 11-octadecenyl acetate | 310.52 | 1.96E-03 | 3 |
| 027519-02-4 | (Z)-9-tricosene | (Z)-9-tricosene | 322.62 | 8.45E-03 | 3 |

